# A hyperglycosylated form of Kv_1.2_ upregulated in LGI1 knockout mice

**DOI:** 10.1101/2025.10.29.685130

**Authors:** Jorge Ramirez-Franco, Marion Sangiardi, Kévin Debreux, Maya Belghazi, Christian Lévêque, Michael Seagar, Oussama El Far

## Abstract

Kv_1_ voltage-gated potassium channels determine key functional neuronal properties. Their activity is modulated by subunit composition and post-translational modifications such as phosphorylation and glycosylation. Using an antibody directed against a phosphotyrosine (Y_458_) located in the C-terminal tail of Kv_1.2_, we identified yet unreported high molecular weight forms of Kv_1.2_ among them, a phosphorylated and heavily glycosylated 100 kDa form. Owing to the significant downregulation of Kv_1.2_ in LGI1-dependent autosomal dominant lateral temporal lobe epilepsy, we investigated, in total brain and the hippocampal formation of both WT and *Lgi1^-/-^* mice, the distribution of phosphoY_458_ Kv_1.2_ and we compared their respective proteomic interactomes with those of Kv_1.2_. In addition to major differences between the interactomes of pY_458_Kv_1.2_ and Kv_1.2_ in WT and *Lgi1^-/-^*, we found a major reshaping of pY_458_ Kv_1.2_ molecular neighbourhood between WT and *Lgi1^-/-^* as well as a significant upregulation of the glycosylated form in *Lgi1^-/-^*.

## Introduction

Disruption of LGI1 function, whether genetic or autoimmune, results in ADLTE and LGI1-associated limbic encephalitis [1] [2]. It is associated with a marked reduction of both total and plasma membrane levels of the delayed rectifying Kv_1_ channels [1] [3] [4] [5]. This decrease is a major contributor to the increased neuronal excitability, and thus presumably to associated epileptic condition. While LGI1 is known to be part of the Kv1-associated proteome [6] [7] [8] [9], the mechanism by which the loss of this extracellular glycoprotein results in Kv_1_ downregulation remains unclear and it can be hypothesised that LGI1 influences the subcellular environment as well as the Kv1 associated-proteome. Ras homolog family member A (RhoA) activity can be linked to receptor tyrosine kinase activity [10] and has been shown to be highly increased by LGI1 knockout [11]. LGI1 may also antagonize NgR1-TROY-RhoA signalling pathway, leading to decreased RhoA activation [11].

The Kv_1_ channels are low voltage-activated K^+^ channels involved in the regulation of neuronal excitability by controlling firing patterns and action-potential (AP) thresholds [12] [13] [14]. They are formed by a homomeric or heteromeric co-assembly of four alpha subunits with auxiliary subunits [15]. D-type α-dendrotoxin-sensitive currents are generated by channels with Kv_1.1_, Kv_1.2_ and Kv_1.6_ subunits and native Kv_1.2_ subunits are present in axonal and synaptic compartments [16] [17] as well as at axonal initial segments [18] [1].

Mutations in these subunits have been shown to be involved in several neurological disorders [19] [20] [21] [22] and functional plasticity in neurons involves a dynamic change in the subcellular distribution of potassium channels [15].

Targeting of Kv_1_ to the membrane, as well as subcellular compartments is complex and heterogeneous. Distinct molecular partners are involved in Kv_1_ targeting and clustering depending on the neuronal compartment concerned. Phosphorylation of Y_458_ in Kv_1.2_ plays a negative role in axonal targeting of Kv_1.2_ and the Y_458_A mutation was shown to increase axonal targeting [17]. The phosphorylation of this tyrosine is a target for physiological regulation by GPCR-coupled mechanisms since Y_458_ phosphorylation is induced by the muscarinic M1 acetylcholine receptor activation and regulates the interaction of Kv_1.2_ with the actin-binding protein cortactin [23]. Although Kv_1_-associated partners Caspr2 and TAG1 are present at the neuronal axonal initial segment (AIS), these proteins as well as DLG4 [24] [6] and ADAM22 [6] are dispensable for Kv1 recruitment in this compartment [16] [25], where other molecular organisers such as DLG2 take over this task [24] [16] [6]. More recently and despite the absence of SCRIB from the proteome of Kv_1.2_ [26], a direct association of Kv_1.2_ with SCRIB was suggested to take place in the AIS and together with PSD93 modulate Kv1 clustering [27]. In addition to the importance of tyrosine phosphorylation in Kv_1.2_ clustering, tyrosine kinase-dependent endocytosis of Kv_1.2_ has been reported to be a mechanism for Kv_1.2_ downregulation [28].

In order to address the potential importance of Y_458_ phosphorylation in determining the molecular environment of Kv_1.2_ and therefore gain insight into its importance in the life cycle of Kv_1.2_ interactions, we developed a phosphoY_458_Kv_1.2_ (pY_458_Kv_1.2_) antibody and, after immunoprecipitation, compared by mass spectrometry the proteomes of pY_458_Kv_1.2_ with the global Kv_1.2_ proteome that we have previously described [26]. Furthermore, since Kv_1.2_ is massively downregulated in *Lgi1^-/-^* mice [1] [26], we addressed the differences in the pY_458_Kv_1.2_ expression and associated proteome between WT and *Lgi1^-/-^* backgrounds.

## Materials and Methods

### Antibodies and reagents

A rabbit polyclonal antibody (ESM_1a) recognizing specifically the phosphoY_458_ (pY_458_) in the C-terminal peptide (aa 454-SKSD(pY)MEIQEGVNNS-468) of mouse Kv_1.2_ was produced. Antibodies recognizing the non-phosphorylated peptide were depleted on an identical but non-phosphorylated peptide column and specific pY_458_ antibodies were affinity purified against the pY_458_Kv_1.2_ peptide (GeneCust). This antibody was used at 5 µg/ml for western blot analysis and 5 µg were used for immunoprecipitation. Kv_1.1_ (K36/15) and Kv_1.2_ (K14/16) antibodies were from Neuromab and used at 1 μg/μl for western blots and immunohistofluorescence experiments and 5 μg for immunoprecipitation. Sumo2/3 antibody (4G11E9) was from GenScript biotech and Ubiquitin antibody (sc-166553) from SantaCruz biotechnology, Inc. HRP-coupled secondary antibodies were from Jackson ImmunoResearch. Unless stated otherwise, chemicals were from Sigma-Aldrich, PNGase and calf intestinal phosphase were from New England Biolabs.

### Dot blots

Phosphorylated and non-phosphorylated peptides (5 ng / dot) were dotted onto a nitrocellulose membrane and classical western blot procedure was used to test the recognition of peptides with anti-Kv_1.2_ and anti-pY_458_Kv_1.2_ peptide antibodies (0.25 µg/ml).

### Western blots

Mouse brains were homogenized in 25 mM Tris-HCl pH 7.4, 150 mM NaCl in the presence of protease and phosphatase inhibitors. Homogenates were recovered in supernatants of a 800×g centrifugation. 40-60 μg of protein were resolved by SDS/PAGE and processed for Western blotting using classical procedures with the indicated antibodies.

### PNGase treatment

After immunoprecipitation, samples were washed and treated with PNGase following the manufacturer instructions. Briefly, solubilized brain extracts from *Lgi1^-/-^* were immunoprecipitated by the pY_458_Kv_1.2_ antibody. Immunoprecipitated material was denatured at 100°C for 10 min and divided in multiple identical aliquots. All aliquots were treated exactly the same way in the presence or absence of PNGase (1µl /20 µg of proteins), incubated for 1h at 37°C and analysed by Western blot. Experiments were performed in triplicates.

### Identification by mass spectrometry of high molecular weight forms of Kv_1.2_

Brain homogenates were solubilized in Tris 25 mM pH 7.4, NaCl 150 mM CHAPS 1% in the presence of phosphatase and protease inhibitors. 4 mg were subjected to immunoprecipitation by 40 µg of polyclonal Kv_1.2_ antibodies. Immunoprecipitated material was denatured at 55°C for 15 min and resolved by electrophoresis on 8% polyacrylamide gel. The migration was stopped when the molecular weight marker of 55 kDa migrated out of the gel. The gel lane was then sliced into 9 distinct pieces along the migration axis and the presence of Kv_1.2_ in each slice was verified by mass spectrometry. Gel bands were processed as previously described [26]. Briefly, gel slices were reduced with dithiothreitol, alkylated with iodoacetamide and digested overnight at 37° in 25 mM ammonium bicarbonate buffer pH 7.4 using Trypsin/Lys-C mix (Promega, Madison, USA). Peptides were extracted three times using 50% acetonitrile (v/v) in water containing 0.1% (v/v) formic acid and dried in a SpeedVac concentrator. Samples were reconstituted in loading buffer (2% acetonitrile (v/v) in water containing 0.1% (v/v) trifluoroacetic acid) and analyzed by LC-MS/MS (nanoHPLC Neo Vanquish coupled to Qexactive Plus from Thermo Fisher Scientific, San Jose, USA). Peptides were first loaded onto a trap column (PepMap Neo 300 µm x 5mm, C18 5µm, 100Ǻ) then separated on an analytical column (EASY-spray PepMap Neo, 75µm x 500 mm, C18 2µm, 100Ǻ) using the following gradient: from 2% to 25% of mobile phase B (20% water, 80% acetonitrile/0.1% formic acid) in A (0.1% formic acid in water) over 90 min, then to 50% B over 20 min. The mass spectrometer was operated in Data Dependant Acquisition positive mode. Full MS scans were acquired at 70 000 resolution (m/z range 350-1900, Auto Gain Control target 3×106) followed by MSMS scans of the top 10 most intense ions (17 500 resolution, isolation window 2m/z, Auto Gain Control 1×105, normalized collision energy 27). Data were searched against the Mus musculus Uniprot database (TaxID=10090, v2025-02-05) using Proteome Discoverer 3.0 software (Thermo Fisher Scientific, San Jose, USA). The mass spectrometry proteomics data have been deposited to the ProteomeXchange consortium via the PRIDE [29] partner repository with the data set identifier PXD069032.

### Biochemical sample preparation, immunoprecipitations and mass spectrometry of pKv_1.2_ proteome

Sample preparation, immunoprecipitation as well as mass spectrometry were performed as in [26]. All samples were treated in triplicates.

Protein quantification was based on the exponentially modified Protein Abundance Index (emPAI) obtained for each identified protein. Three biological samples from WT and *Lgi1^-/-^* were immunoprecipitated using pY_458_Kv_1.2_ antibody. Two parallel biological samples from each genotype were immunoprecipitated using control rabbit antibodies (NIAB). Data were analyzed and for each protein in the partners lists, the emPAI values from the NIAB samples were averaged; if a protein was detected in only one NIAB sample, the missing value was treated as zero before averaging resulting in: *Avg emPAI(NIAB) =* [*emPAI(NIAB 1) + emPAI(NIAB2)*] */ 2 or Avg emPAI(NIAB) =* [*emPAI(NIAB present) + 0*] */ 2*

For each pY_458_Kv_1.2_ antibody sample, the average NIAB emPAI value for a given protein was subtracted to yield a background-corrected emPAI value (corrected emPAI). Subsequently, the ratio between the pY_458_Kv_1.2_ antibody and NIAB emPAI values was calculated for each protein: *emPAI Ratio* = *emPAI pY_458_Kv_1.2_ / Avg emPAI NIAB*. Proteins were retained for further analysis only if they met all of the following criteria: (i) Ratio ≥ 3, (ii) number of unique peptide sequences > 1, (iii) presence in more than one pY_458_Kv_1.2_ antibody sample, and (iv) corrected emPAI > 0.1.

To remove highly abundant or nonspecific proteins, a semantic filter was applied. Proteins whose descriptions contained any of the following terms were excluded from the analysis: “Complement,” “DNA binding protein”, “Initiation,” “Elongation,” “Haemoglobin,” “Immunoglobulin,” “Keratin”, “Mitochondrial”, “Nuclear,” “RNA,” “RNA splicing”, “exonuclease”, “Ribosomal,” “Transcription,” “Tubulin,” or “Transcriptional.” Proteins that passed both the quantitative and semantic filters were retained, and their corrected emPAI values were averaged across replicates for further analysis. Candidate proteins were classified using the Genecards and pantherdb web sites (www.genecards.org; www.pantherdb.org) and association networks were analysed using the String protein server (https://string-db.org).

### Immunohistofluorescence staining of fixed brains

All experiments were performed in accordance with the European and institutional guidelines for the care and use of laboratory animals (Council Directive 86/609/EEC and French National Research Council) and approved by the local authority (Préfecture des Bouches-du-Rhône, Marseille). Fixed brains of P14-P16 C57BL/6 wild-type mice or *Lgi1*^-/-^ littermates of either sex were sliced and stained as described in [9]. Three animals per genotype were used for each of the immunohistofluorescence conditions tested.

### Image analysis

For quantification of immunolabeling differences in WT and *Lgi1*^-/-^ animals, two different strategies were used. In panel C of Figure 5, three straight lines were placed along the different CA3 strata in each field of view (n=3 animals, n=6 slices). Fluorescent values were collected as arbitrary fluorescent units (afu) and normalized to the first point of the WT traces. For the panels 5D and 5E, 5 random ROIs (Regions of Interest) where placed in each of the CA3 hippocampal strata (n=3 animals, n=6 Slices). Fluorescence values where collected and normalized to the average fluorescence value of WT in each of the strata. For co-localization analysis, the Just Another Colocalization Plugin (JACOP) was used in its BIOP version [30] and the Persons’ Correlation Coefficients (PCC) were computed over z-stacks consisting of 10 slices considering z-slices separately but averaging values in a single data point.

## Results

### Identification of phosphorylated high molecular weight forms of **Kv_1.2_**

We generated a polyclonal antibody recognizing the phosphorylated form of Kv_1.2_ on Y_458_ (pY_458_Kv_1.2_). The specificity of this antibodies was first addressed by dot blot. As shown in ESM_1a, pY_458_Kv_1.2_ antibodies recognize the phosphorylated peptide immunogen but not the very same immunogen in its non-phosphorylated state. Of note, our previously reported antibody, generated against the non-phosphorylated peptide [26], cross-reacts with the phosphorylated peptide in dot blots and therefore recognise both peptides (ESM_1a). Sequence alignment of Kv_1_ C-terminal sequences shows a significant similarity in the Y_458_ environment between Kv_1.1_, Kv_1.2_, Kv_1.3_ and Kv_1.4_, (ESM_1b) and suggests that the pY_458_Kv_1.2_ antibody may cross-react with other phosphorylated Kv_1_ subunits. Immunoprecipitation experiments followed by mass spectrometry shows that pY_458_Kv_1.2_ antibody captures only Kv_1.1_ and Kv_1.2_ from WT mice and the total absence of other Kv_1_ subunits (ESM_1c) that were otherwise associated with Kv_1.2_ [26] namely Kv_1.3_, Kv_1.4_, Kv_1.5_ and Kv_1_._6_ subunits. These data argue against a potential cross-reactivity of the pY_458_Kv_1.2_ antibodies with other Kv_1_ alpha subunits than Kv_1.1_. Western blot analysis of brain homogenates (ESM_2) shows that pY_458_Kv_1.2_ antibody identifies, in addition to the circa 70 kDa principal band, 3 bands of higher molecular weight (circa 100, 130 and >200 kDa) that are not recognized in homogenates by the K14/16 Kv_1.2_ Neuromab Kv_1.2_ antibody as well as our previously characterized polyclonal Kv_1.2_ antibody [26]. In order to confirm the identity of Kv_1.2_ reactive species at the observed high molecular weights, Kv_1.2_ was immunoprecipitated with the polyclonal Kv_1.2_ antibody, that, as shown in ESM_1a, recognizes both phosphorylated and non-phosphorylated peptides, and subjected to polyacrylamide gel electrophoresis. Slices at different migration positions of the migration lane were analysed by mass spectrometry for the presence or absence of Kv_1.2_. As shown in Fig. 1a, Kv_1.2_ peptides were found at distinct migration levels corresponding to the observed high molecular weight forms of Kv_1.2_. Since Kv_1.2_ is massively decreased in *Lgi1^-/-^* we investigated potential modification in the expression of pY_458_Kv_1.2_ in *Lgi1^-/-^*. Similar to the previously described expression decrease in the 70 kDa form of Kv_1.2_ in *Lgi1^-/-^*, the a=70 kDa pY_458_Kv_1.2_ also decrease by at least two-fold (ESM_2). Two of the newly detected protein bands did not vary (c=130 & d=>200 kDa) and one (b=100 kDa) increased by > 70% in *Lgi1^-/-^* (ESM_2) with b/a ratio of 1.68 ± 1.8 in WT and 3.26 ± 0.66 in *Lgi1^-/-^*.

**Fig. 1.**
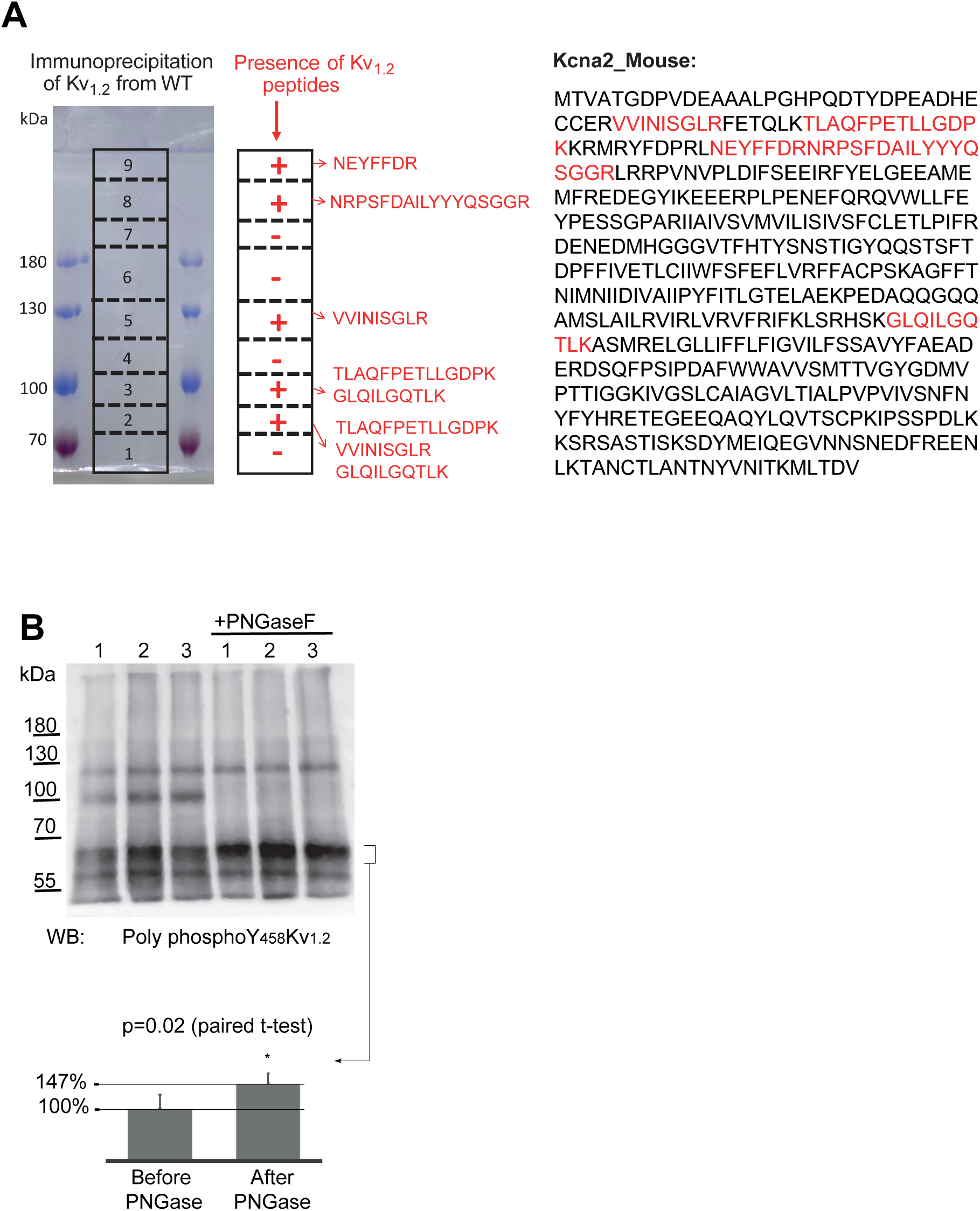
Identification and glycosylation profile of the molecular entities immunoprecipitated by pY_458_Kv_1.2_. **a** Kv_1.2_ was immunoprecipitated by the polydonal Kv_1.2_ antibody from WT brain extracts and subjected to SDS PAGE. Mass spectrometry analysis was performed on the indicated polyacrylamide gel slices and the presence or absence of Kv_1.2_ peptides is reported (in red). The position of the found peptides in the linear Kv_1.2_ sequence is indicated in the Kv_1.2_ sequence (right side). **b,** Kv_1.2_ was immunoprecipitated by the polydonal Kv_1.2_ and the immunoprecipitated material was subjected or not to PNGase treatment. Samples were analysed by Western blot with the pY_458_Kv_1.2_. The circa 70 kDa bands were quantified and their intensity plotted in histograms. p values (paired t-test) are reported under each Western blot (n=3)

The large increase in the molecular weight of Kv_1.2_ is unlikely due to phosphorylation and may indicate additional post-translational modifications such as ubiquitination, sumoylation or glycosylation. An increase in the apparent molecular weight of Kv_1.3_ has been reported upon ubiquitination [31] and a highly glycosylated form of Kv_1.2_ with a large increase in the apparent molecular weight was previously reported in a heterologous system [32]. None of the Kv_1.2_ high molecular weight bands recognized and enriched by the pY_458_Kv_1.2_ antibody was recognized by ubiquitin or Sumo2/3 antibodies (ESM_3). However, the 100 kDa band completely disappeared after treatment with PNGaseF while the 70 kDa band increased (Fig. 1b). This indicates that the 100 kDa species corresponds to a highly glycosylated form of Kv_1.2_. All these data indicate that the antibody raised against the phosphoY_458_ in the C-terminal domain of Kv_1.2_ recognizes a phosphorylated form of the well characterized 70 kDa but also other high molecular weight forms with potentially differential post-translational modifications.

### Distribution of pY_458_Kv_1.2_ in WT versus *Lgi1^-/-^*hippocampal regions

In order to address the localisation of pY_458_Kv_1.2_ and its distribution throughout the brain and the hippocampal formation, we performed immunostainings on whole brain slices of WT animals. The staining pattern of the pY_458_Kv_1.2_ antibody revealed an intense labeling of myelinated forebrain and midbrain axonal tracts (ESM_ **4**). Moreover, a more diffuse yet clearly detectable labeling pattern was observed in several hippocampal regions, particularly within the stratum lucidum (SL) of CA3 (ESM_4 and Figs. 2a, 2b, and 2c). In order to address the differences in the distribution of pY_458_Kv_1.2_ in WT and *Lgi1^-/-^*, we performed a comparative analysis of the staining pattern in WT vs *Lgi1^-/-^* hippocampal slices (WT n=3 animals, *Lgi1^-/-^* n=3 animals). As shown in Fig. 2 (Figs. 2a, 2b, and 2c), WT hippocampal slices show weak staining, whereas *Lgi1^⁻/⁻^* slices exhibit intense labeling across all analyzed CA3 hippocampal strata (SO: Stratum Oriens; SL: Stratum Lucidum; SR: Stratum Radiatum; Figs. 2c and 6d; n = 3 animals per genotype). We focused our analysis on CA3 due to its enriched expression of LGI1 and the reported selective reduction of Kv_1_-family channel levels in this region in *Lgi1⁻^/^⁻* mice [1] [26]. Normalized fluorescence values (Fig. 2e) confirmed this observation, with significantly increased signals in *Lgi1^⁻/⁻^* compared to WT littermates in all layers of CA3: SO (1.55 ± 0.04 vs. 1.00 ± 0.07), SL (2.17 ± 0.05 vs. 1.00 ± 0.07), and SR (2.04 ± 0.07 vs. 1.00 ± 0.07). In order to ascertain whether this increase in pY_458_Kv_1.2_ immunoreactivity was of presynaptic or postsynaptic origin, we performed double immunostaining against pY_458_Kv_1.2_ and the presynaptic marker synaptophysin (n=2 animals, Fig. 2f). Co-localization analyses yielded very low and similar Pearson correlation coefficients between the pY_458_Kv_1.2_ signal and synaptophysin in both genotypes (PCC=0.0199 ± 0.0185 in WT and PCC=0.0263 ± 0.0146 in *Lgi1^⁻/⁻^* mice), indicating that the hippocampal pY_458_Kv_1.2_ signal was mostly of non-presynaptic origin. This large increase in pY_458_Kv_1.2_ staining in *Lgi1^-/-^* background may reflect the observed increase in the intensity of the glycosylated 100 kDa form of Kv_1.2_ by western blot but potentially also a differential accessibility of the anti pY_458_Kv_1.2_ epitope between WT and *Lgi1^-/-^*.

**Fig. 2.**
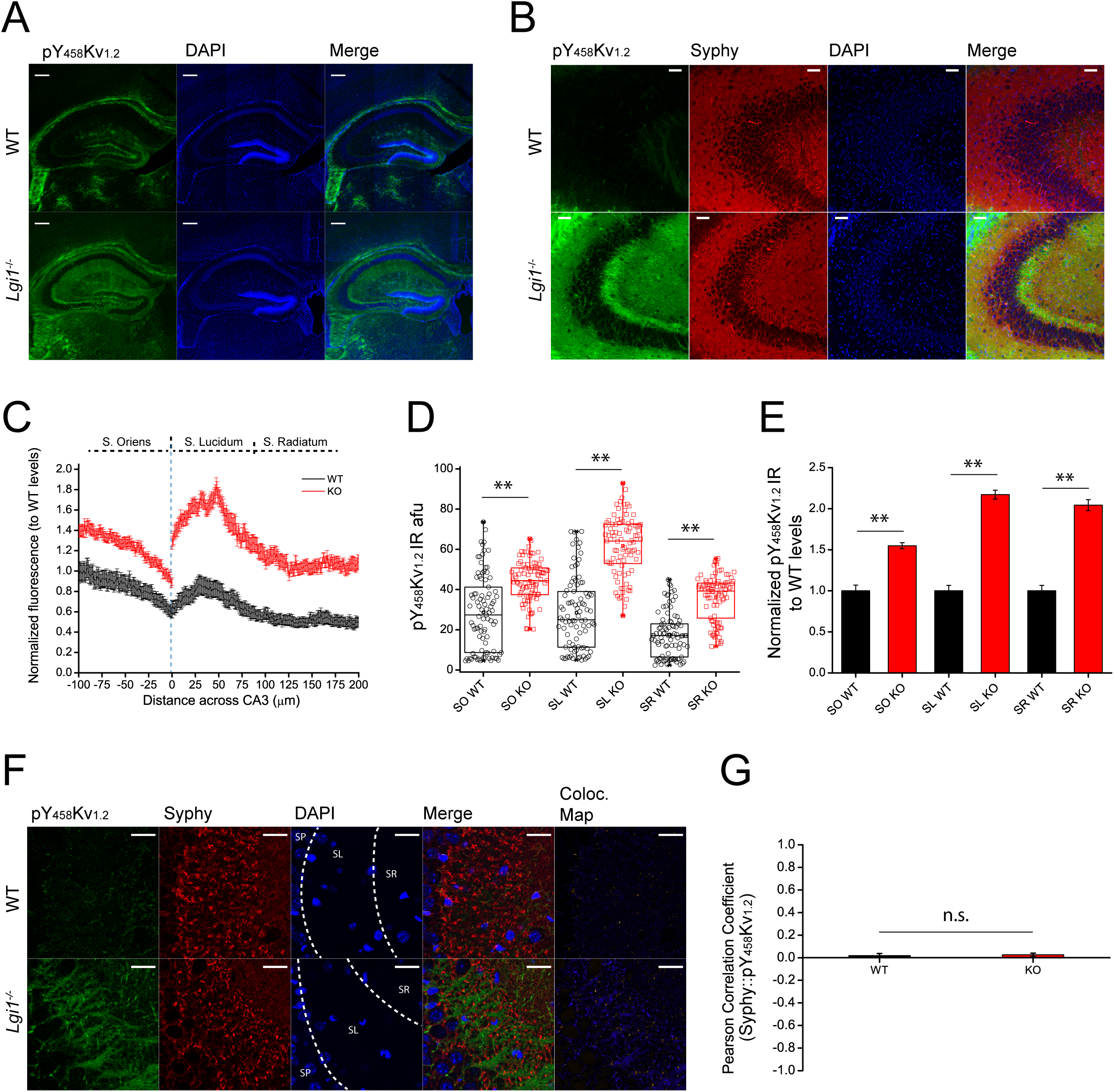
lmmunofluorescence distribution of pY_458_Kv_1.2_ in the hippocampal formation of WT and *Lgi1^-/-^*. **a** Full hippocampal immunostaining with anti-pY_458_Kv_1.2_ in WT (top) and *Lgi1^-/-^* (bottom) showing pY_458_Kv_1.2_ immunoreactivity (green), DAPI (blue), and merged images. Scale bar = 250 µm. **b** Detailed views of CA3 hippocampal regions showing immunolabeling against pY_458_Kv_1.2_ (green), Synaptophysin (red), DAPI (blue), and merged images. Scale bar = 50 µm. c Average line trace quantification of immunolabeling shown as normalized signal of arbitrary fluorescence units (afu). **N** = 3 animals, n = 6 slices/animal; 3 lines per slice. Mean ± SEM. **d** Box plots of randomly placed ROls across CA3 hippocampal strata in WT (black dots and boxes) and *Lgi1_-/-_(red* dots and boxes). **N** = 3 animals; n = 6 slices/animal; 5 ROls per slice and stratum. (SO WT = 28.66 ± 1.97 vs *Lgi1^-/-^*= 44.40 ± 1.04, p = 3.75 x 10^-44^; SL WT = 28.46 ± 1.87 vs *Lgi1^-/-^* = 61.79 ± 1.54, p = 1.38 x 10^-44^; SL WT = 17.60 ± 1.17 vs *Lgi1^-/-^* = 35.96 ± 1.18, p = 1.08 x 10^-15^). p < 0.01, One-way ANOVA followed by Bonferroni’s test for mean comparisons. e Bar graph comparisons of the normalized values shown in panel C for WT (black bars) and *Lgi1^-/-^* (red bars). Data are Mean± SEM: (SO WT= 1.00 ± 0.07 vs *Lgi1^-/-^* = 1.55 ± 0.04, p = 5.41 x 10^-s^; SL WT= 1.00 ± 0.07 vs *Lgi1^-/-^* = 2.17 ± 0.05, p = 2.66 x 10^-35^; SR WT= 1.00 ± 0.07 vs *Lgi1^-/-^* = 2.04 ± 0.07, p = 6.55 x 10^-29^). p < 0.01, One-way ANOVA followed by Bonferroni’s test for mean comparisons. f Detailed views of the stratum lucidum of CA3 in WT (top) and *Lgi1^-/-^* (bottom) showing staining for anti-pY_458_Kv_1.2_ (green), anti­ Synaptophysin (red), DAPI (blue), and merged images. Right-most panels show a colocalization map using an Intensity Correlation Analysis (ICA) Lookup Table (LUT). Scale bar = 25 µm. g Pearson’s correlation coefficients (PCC) for co-localization of pY_458_Kv_1.2_ and Synaptophysin in WT (black bars) and *Lgi1^-/-^* (red bars). N = 2 animals, n = 4 slices/animal. Data are Mean ± SEM. Two-sample t-test; n.s., non-significant (WT= 0.0199 ± 0.0185 vs *Lgi1^-/-^* = 0.0263 ± 0.0146, p = 0.79).

### Subunit composition of Kv_1_ channels containing pY_458_Kv_1.2_

Mass spectrometry analysis of immunoprecipitated samples shows an association between Kv_1.2,_ Kv_1.1_ and Kvβ_2_ subunits in WT samples. In contrast, neither Kv_1.1_ nor Kvβ_2_ were recovered in *Lgi1^-/-^* (ESM_1c). Of note, the recovery yield with anti-pY_458_Kv_1.2_ (emPAI = 0.36 in WT; emPAI = 0.14 in *Lgi1^-/-^*) is much smaller than with anti-Kv_1.2_ (emPAI = 2.9 in WT and 2.8 in *Lgi1^-/-^*) [26]. This difference cannot be attributed to a limited amount of material upon immunoprecipitation since we used an excess of solubilized WT or *Lgi1^-/-^* brain extracts. This indicates thus that pY_458_Kv_1.2_ is only a minor component of the entire Kv_1.2_ population. Also, while the association of Kvβ_1_ & Kvβ_2_ with Kv_1.2_ is very consequent (Kvβ emPAI > Kvα emPAI) [26] (ESM_1), the association of Kvβ_2_ with pY_458_Kv_1.2_ is clearly lower with Kvα emPAI > Kvβ emPAI).

### Comparison of Kv_1.2_ and pY_458_Kv_1.2_ proteomes in WT and *Lgi1^-/--^*

Among the partners that were exclusive to Kv_1.2_ or pY_458_Kv_1.2_ in WT background compared to *Lgi1^-/-^*, none were found common (Fig. 3a). Among those that were exclusive to Kv_1.2_ or pY_458_Kv_1.2_ in *Lgi1*^-/-^ compared to WT, only four showed to be common (Fig. 3b). Only seven partners were shared between the common partner lists of Kv_1.2_ and pY_458_Kv_1.2_ across both genotypes, (Fig. 3c).

**Fig. 3.**
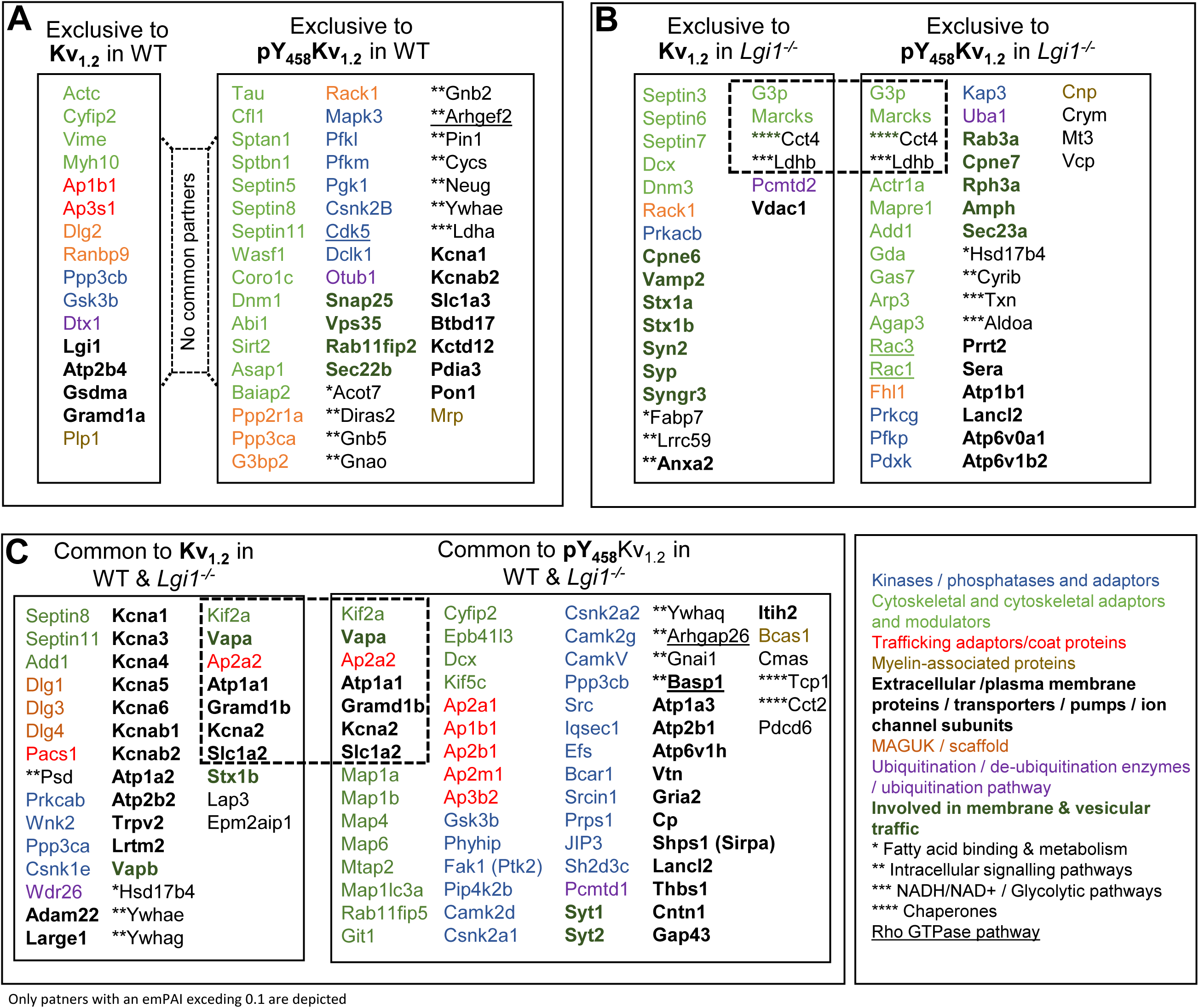
Comparative listing of Kv_1.2_ versus pY_458_Kv_1.2_ partners in WT and *Lgi^-/-^*. **a** In WT. **b** In *Lgit-1^-/-^*. **c** common partners in both genotypes

#### Association with cytoskeletal elements

We previously showed [26] that LGI1 modulates Kv_1.2_ association with the actin network since, in the absence of LGI1, Kv_1.2_ loses interaction with cytoskeletal actin but remains associated with adducin and doublecortin (Fig. 3a, b). The present data show that in WT, pY_458_Kv_1.2_ is not associated with cytoskeletal actin but with actin related protein 1a (actr1a) that is a component of the dynactin complex involved in microtubule based intracellular transport. It is also associated with the spectrin / tau / microtubule network [33]. The presence of actr1a, spectrin as well as cofilin and other actin regulating proteins in association with pY_458_Kv_1.2_ (Fig. 3a; Fig. 4) in WT but not in *Lgi1*^-/-^ may indicate a link between pY_458_Kv_1.2_ and endocytic pathways [34] and highlights an important difference in the fate of Kv_1.2_ relative to pY_458_Kv_1.2_ as well as pY_458_Kv_1.2_ in WT and *Lgi1*^-/-^. Interestingly, the association of pY_458_Kv_1.2_ with the microtubule associated tether protein VAPA [35] and the non-vesicular lipid transfer protein ASTRB [36] [37] is strongly decreased in *Lgi1^-/-^*, pointing towards a mislocalization of pY_458_Kv_1.2_ in *Lgi1^-/-^*.

**Fig. 4.**
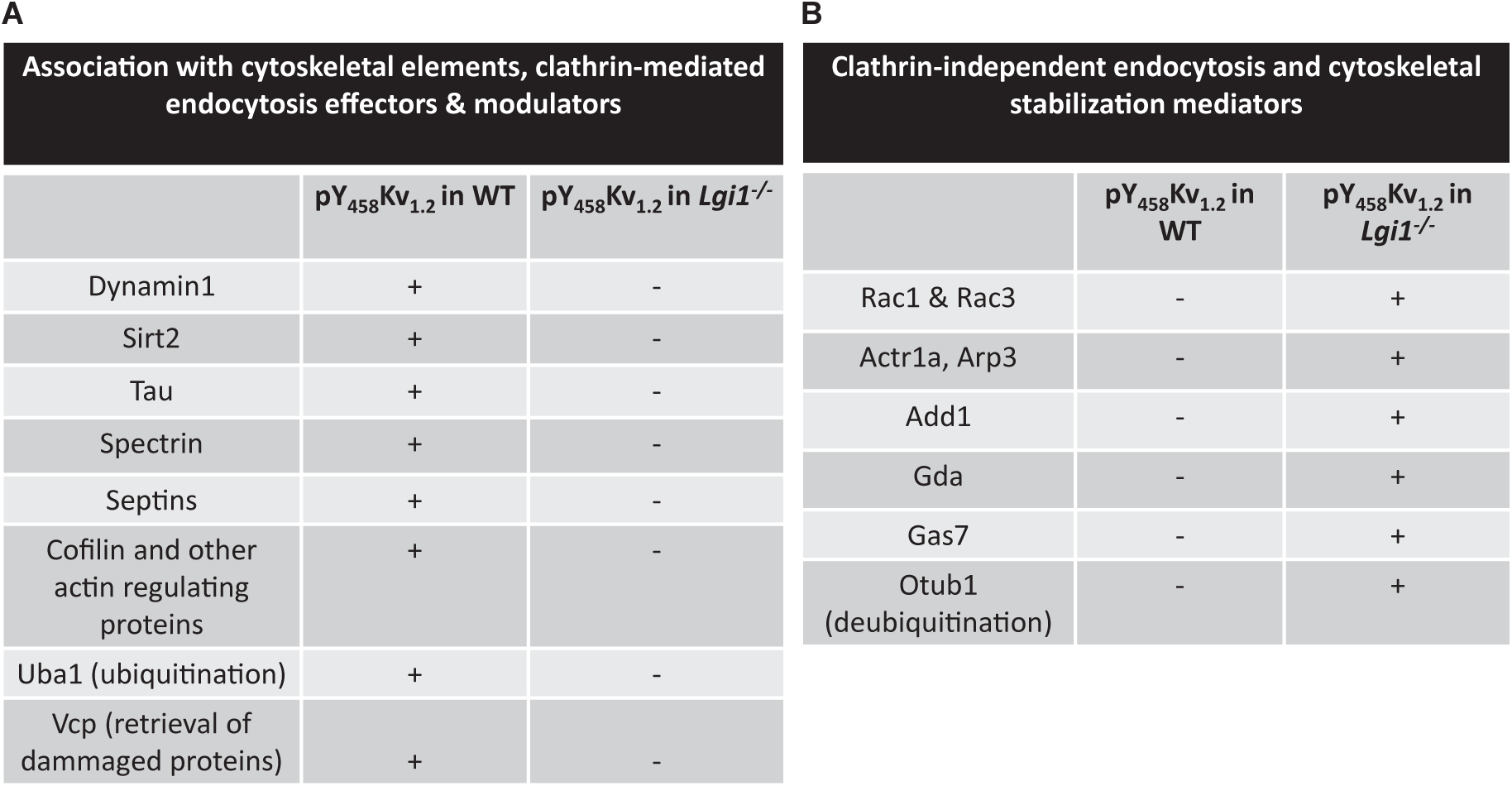
Comparative listing of cytoskeletal elements and endocytotic effectors associated with pY_458_Kv_1.2_ in WT and *Lgi1^-/-^*. **a, b** report cytoskeletal links and Clathrin-mediated versus Clathrin-independent endocytotic effectors respectively

Comparison of pY_458_Kv_1.2_ proteome between WT and *Lgi1^-/-^* shows that, pY_458_Kv_1.2_ in WT is associated with indicators of the clathrin-dependent endocytic pathway (Fig. 4a), while pY_458_Kv_1.2_ in *Lgi1^-/-^* seems to be handled differently (Fig. 4a). Upon comparison of all Kv_1.2_ and pY_458_Kv_1.2_ partners, 15 and 16 partners were common to WT and *Lgi1^-/-^* respectively (ESM_5). Among these, 7 were common in both WT and *Lgi1^-/-^*. Analysis of the Go terms of non-common partners show a particular enrichment of the Go term “cytoskeleton” for pY_458_Kv_1.2_ partners in *Lgi1^-/-^* (ESM_5) suggesting that despite the absence of interaction with the actin and spectrin, pY_458_Kv_1.2_ in *Lgi1^-/-^* may be subject to a differential regulation of interaction with cytoskeletal links. A closer look at exclusive pY_458_Kv_1.2_ partners in *Lgi1^-/-^* (Fig. 4b) shows the association with clathrin-independent endocytosis adaptors and cytoskeletal stabilization mediators (Arp3, Actr1a, Rac1 & 3, Add1, Gda, Gas7 and Otub1) suggesting a particular compensatory mechanism stabilizing cytoskeletal anchoring. Arp3 is the ATP-binding component of Arp2/3 complex involved in actin polymerization [38]: Actr1a / alpha centractin is a component of a multi-subunit complex involved in microtubule based vesicle motility [39]. Rac1 & 3 are important effectors in actin filaments polymerisation [40]. Add1 is a membrane-cytoskeleton-associated protein promoting the assembly of the spectrin-actin network [41] and Gas7 is an actin-filament binding protein [42]. By its deubiquitination activity, Otub1 modulate the endocytotic fate of certain proteins. Of note, Kv_1.2_ and pY_458_Kv_1.2_ are associated, only in WT, with Cyfip2 that negatively regulates Rac1-driven cytoskeletal remodelling and the actin-binding protein GAS7 highlighting the importance of LGI1 in the association of Kv_1.2_ and pY_458_Kv_1.2_ with the cytoskeletal network.

#### Association with septins

As previously described [26], the association of Kv_1.2_ with septins is different between WT and *Lgi1^-/-^*. In WT, Kv_1.2_ associates with only two septins (8 and 11) while in *Lgi1^-/-^*, three additional septins, the neuron-specific septin 3 implicated in actin filament and microtubule dynamics [43] as well as septins 6 and 7 are also present (Fig. 5a). We find that pY_458_Kv_1.2_ immunoprecipitated from WT background is also associated with septins 8 and 11 like Kv_1.2_ but also with an additional septin (septin 5) (Fig. 5a). Suprisingly, in *Lgi1^-/-^*, pY_458_Kv_1.2_ totally loses association with all septins while its non-phosphorylated equivalent is strongly associated with septins 3, 6, 7, 8 and 11. These data suggest that septins may play a crucial role in shaping the localisation of pY_458_Kv_1.2_ in *Lgi1^-/-^*, especially through an increase in association with Kv_1.2_ and a complete loss of association with pY_458_Kv_1.2_. These data highlight again an important difference in the fate of pY_458_Kv_1.2_ between WT and *Lgi1^-/-^*.

**Fig. 5.**
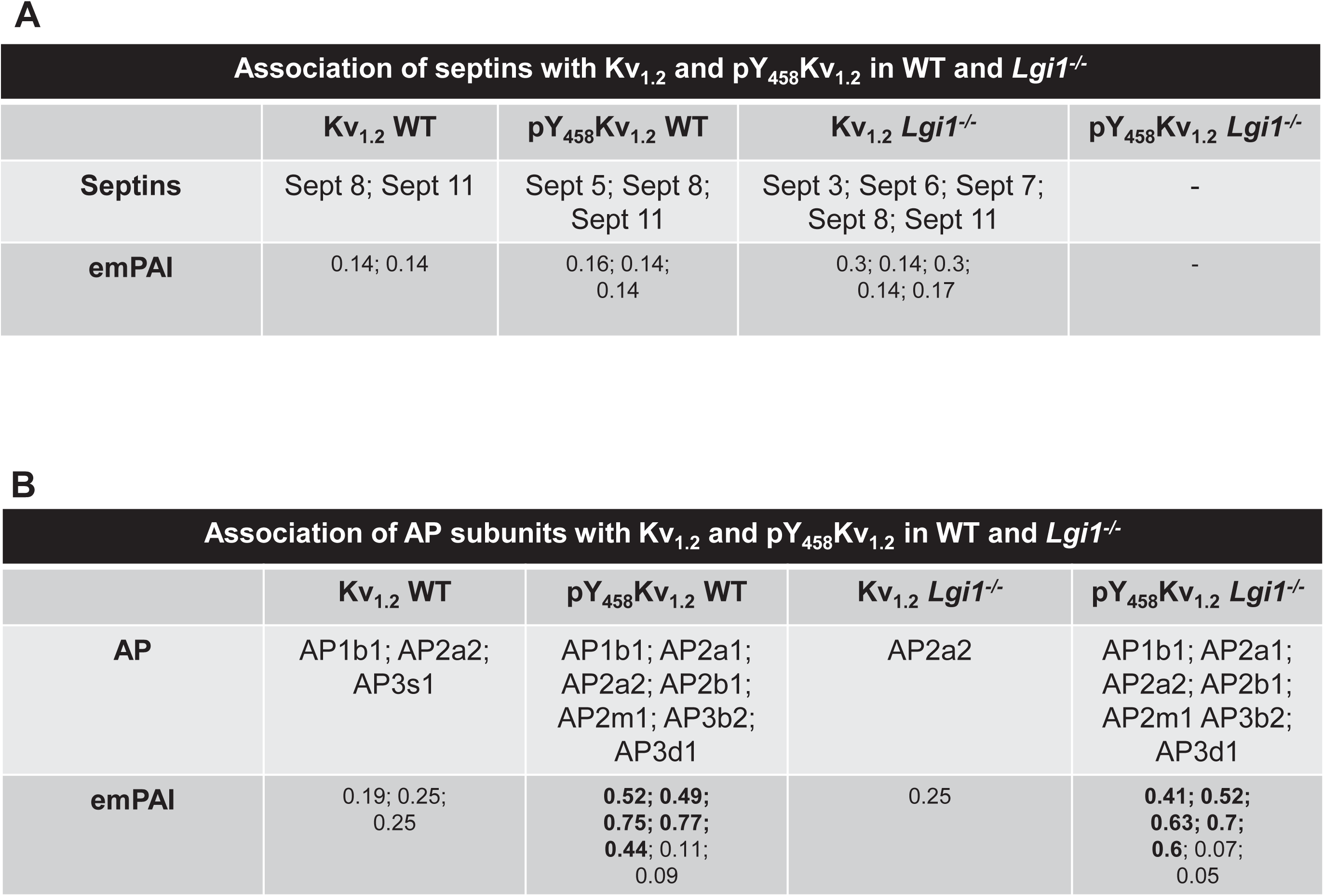
Septins and adaptor proteins associated with Kv_1.2_ and pY_458_Kv_1.2_ in WT and *Lgi1^-/-^*. **a** Comparative listing of Kv_1.2_ and pY_458_Kv_1.2_ association with septins **b** Comparative listing of Kv_1.2_ and pY_458_Kv_1.2_ association with adaptor proteins

#### Association with adaptor protein

Following LGI1 depletion, Kv_1.2_ loses association with AP1 and AP3 subunits but still associates with AP2 via AP2a2 [26] (Fig. 5b).

We find that AP1, AP2 and AP3 subunits [44] (AP3b2; neuron-specific subunit of the AP3 complex, AP3d1, AP1b1, AP2b1, AP2a1, AP2m1) are associated with pY_458_Kv_1.2_ in both genotypes with no fundamental difference between WT and *Lgi1^-/-^* (Fig. 5b).

#### Proteins implicated in LGI1-deficiency associated-pathology

We previously described 42 common molecular partners of Kv_1.2_ in both WT and *Lgi1^-/-^* genetic backgrounds [26] (ESM_6). Interestingly, 35 of these partners are not present in the proteome of pY_458_Kv_1.2_ in particular ADAM22, DLG1, 3,4 as well as 14-3-3 γ and ε, all implicated in LGI1-deficiency associated-pathology.

#### Association with exo-endocytotic proteins

We previously reported that Kv_1.2_ is only weakly associated with exo-endocytic markers in WT background and that the absence of *Lgi1^-/-^* induces stronger association with these markers and with the somatodendritic endocytic marker dynamin 3 [26]. The current analysis shows that the association of Kv_1.2_ and pY_458_Kv_1.2_ with exo-endocytic markers is different already in WT background with pY_458_Kv_1.2_ being significantly associated with exo-endocytic markers (Fig. 6a). The association of pY_458_Kv_1.2_ with dynamin 1 and not dynamin 3 indicate a fundamental difference in the localisation of the major analysed pool of pY_458_Kv_1.2_ in WT suggesting that it is actively endocytosed in presynaptic terminals. pY_458_Kv_1.2_ is also associated with the vacuolar protein sorting ortholog 35 (Vps35) that participates in endosomal sorting. Surprisingly, and despite the enrichment of adaptor proteins with pY_458_Kv_1.2_ in *Lgi1^-/-^*, dynamin1 association is lost indicating potentially either that the endocytic process is hindered or that it takes place in a dynamin-independent pathway. While the ubiquitin removal enzyme OTUB1 which opposes ubiquitination and prevent degradation, is exclusively associated with pY_458_Kv_1.2_ in WT background, the ubiquitin activation E1 enzyme UBA1 is only immunoprecipitated with pY_458_Kv_1.2_ in *Lgi1^-/-^.* In parallel to UBA1, the valosin-containing protein (Vcp) is only found in the proteome of pY_458_Kv_1.2_ in *Lgi1^-/-^.* Vcp interacts with ubiquitinated proteins helping their extraction from membranes or protein complexes before degradation [45]. Together, these observations suggest that in *Lgi1^-/-^,* the retrieval of pY_458_Kv_1.2_ from membranes becomes dynamin independent. The difference in the association with dynamin may also reflect a change in dynamic regulation (exo/endocytosis) of the glycosylated Kv_1.2_ that could account for the plasticity of intrinsic excitability.

**Fig. 6.**
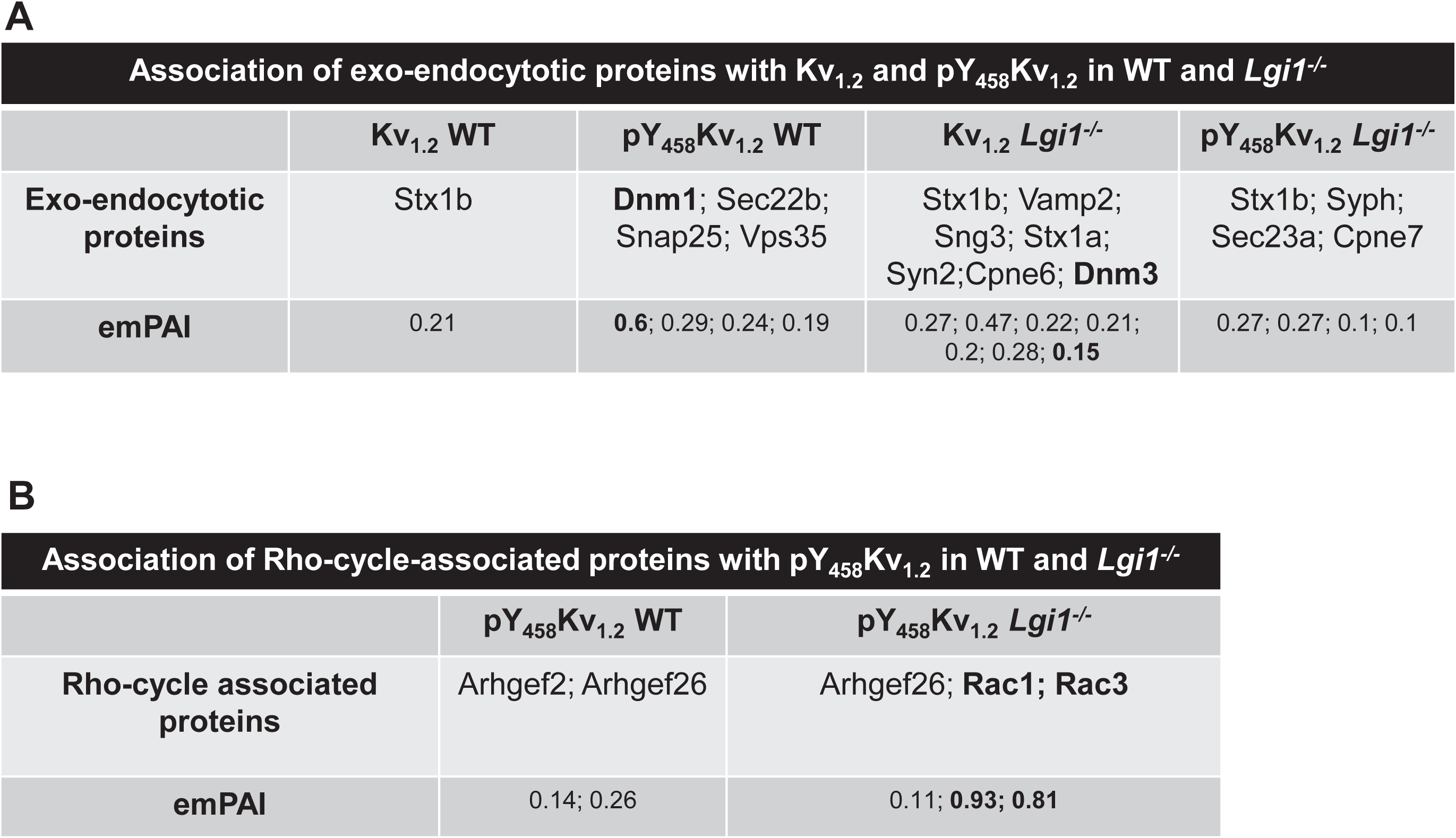
Exo-endocytotic and Rho-cycle-associated proteins associated with pY_458_Kv_1.2_ in WT and *Lgi1^-/-^*. **a** Comparative listing of Kv_1.2_ and pY_458_Kv_1.2_ association with exo-endocytotic proteins, **b** Comparative listing of pY_458_Kv_1.2_ association with Rho-cycle-associated proteins in WT and *Lgi1^-/-^*

#### Rho GTPase-associated pathways with pY_458_Kv_1.2_ in *Lgi1^-/-^*

Guanine nucleotide exchange factors (GEFs) turn on membrane localized small G-proteins signalling by catalysing the exchange of GDP with GTP, GTPase-activating proteins (GAPs) terminate their signalling by inducing GTP hydrolysis [46] [47]. Arhgef2 and Arhgap26 are RhoA effectors implicated in microtubule cytoskeletal dynamics. Upon dissociation from microtubules, Arhgef2 stimulates membrane-bound RhoA leading to actin stress fibre formation that can influence membrane trafficking and endocytosis while Arhgap26 inhibits membrane-bound RhoA activity and stabilises synapses. While we did not observe an association of GEFs and GAPs with Kv_1.2_ [26], the association of pY_458_Kv_1.2_ with the RhoA GAP (Arhgap26) occurs in both WT and *Lgi1^-/-^* genotypes, its association with RhoA GEF (Arhgef2) occurs only in WT background (Fig. 6b). This association again points towards distinct signalling pathways associated with Kv_1.2_ and pY_458_Kv_1.2_ and that LGI1 ablation influences the association of pY_458_Kv_1.2_ with the Rho GEF Arhgef2. This is especially interesting knowing that RhoA activity is significantly increased in *Lgi1^-/-^* [11].

The association of Rac1 and Rac3 with pY_458_Kv_1.2_ occurs only in *Lgi1^-/-^* and with high emPAIs. Rac 3 has been shown to be important in neuronal spine formation [48] and its enrichment in pY_458_Kv_1.2_ in *Lgi1^-/-^* can be related to the decrease in spine pruning observed in in these animals. While Rac 1 is ubiquitous, Rac3 is expressed in neurons and is strongly expressed in CA1-CA3 in the hippocampus [48]. Also, Rac1 and microtubules, are major players in clathrin-independent endocytosis [49]. The enrichment of pY_458_ Kv_1.2_ proteome with microtubules and Rac1 and the association of pY_458_Kv_1.2_ with Rac1 only in *Lgi1^-/-^* again suggest that pY_458_Kv_1.2_ has a different fate between WT and *Lgi1^-/-^* and that in the absence of LGI1, it may be endocytosed through clathrin-independent pathways (Fig. 4b).

#### RACK1 association with Kv_1.2_ and pY_458_Kv_1.2_

In contrast to Kv_1.2_ and independently from the presence or absence of LGI1, pY_458_Kv_1.2_ is associated with a number of kinases and kinase scaffolds (Fig. 3, ESM_7) suggesting that it is associated with an enzymatically active environment and kinase hotspots.

The WD-repeat family member RACK1 (Receptor for activated C Kinase 1) [50] is a shuttling and scaffolding protein. It contributes to several aspects of cellular function and is a key mediator of various pathways. It binds and stabilizes active or inactive conformations of its partners (i.e. stabilizing inactive Src and Fyn kinases and active PKCβII). Interestingly, RACK1 is the only common partner between Kv_1.2_ in *Lgi1^-/-^* and pY_458_Kv_1.2_ in WT. In the list of Kv_1.2_ partners specifically acquired in *Lgi1^-/-^*, we found the cAMP-dependent protein kinase catalytic subunit beta (Prkacb) (Fig. 3b). This association is compatible with the presence of RACK1 among the list of partners and the known involvement of RACK1 in the adenylate cyclase signalling pathway. In contrast, this protein was not a pY_458_Kv_1.2_ partner in WT. Instead, we found a large number of kinases and phosphatases as well as scaffolds and G protein subunits (Fig. 1c) compatible the hub function of RACK1. Among these proteins, the Gα (Gnao) and Gi (Gnb2), Gβ (Gnb5), serine/threonine-protein kinase DCLK1, mitogen-activated protein kinase 3, cyclin-dependent-like kinase 5 Cdk5), serine/threonine-protein phosphatase 2A, serine/threonine-protein phosphatase 2B catalytic subunit alpha, the casein kinase II subunit beta (Csnk2b), Wiskott-Aldrich syndrome protein family member 1 (Wasf1), which is a downstream effector of tyrosine kinase receptors and small GTPases implicated in actin cytoskeleton organization. These data highlight a fundamental difference in the signalling platforms associated with Kv_1.2_ and pY_458_Kv_1.2_ in WT animals. In addition, since Kv_1.2_ joins RACK1 platforms in *Lgi1^-/-^* and pY_458_Kv_1.2_ leaves them, LGI1 appears to be a key factor in organizing signalling platfoms associated with Kv_1.2_ and pY_458_Kv_1.2_ channels.

#### CNPase is specifically associated with pY_458_Kv_1.2_ in *Lgi1^-/-^*

Among the proteins that are specific to pY_458_Kv_1.2_ versus Kv_1.2_ in *Lgi1^-/-^*, we found 2’,3’-cyclic-nucleotide 3’-phosphodiesterase (CNPase/Cnp or CN37) (Fig. 3b). Cnp is the most abundant protein in myelin [51] and is a potential auto-antigen in multiple sclerosis [52]. Variability in its expression levels has been linked to major depressive disorder [53] [54] and schizophrenia [55] [56].

#### Association of pY_458_Kv_1.2_ with microtubule-associated proteins

A specific characteristic of pY_458_Kv_1.2_ is its association with microtubule-associated proteins (Fig. 3c). MAP1b, MAP1 light chain 3a, MAP1a, MAP6, and MAP4 (respectively from the most to the least abundant) were exclusively immunoprecipitated with pY_458_Kv_1.2_ from both WT and *Lgi1^-/-^* however, pY_458_Kv_1.2_ from *Lgi1^-/-^* was significantly less associated with MAPs (ESM_7). The glycogen synthase kinase-3 beta (GSK3b) that controls the activity of MAP1a and b [57] [58] was also recovered although equally enriched from WT and *Lgi1^-/-^* backgrounds. These data suggest that, in *Lgi1^-/-^*, pY_458_Kv_1.2_ may be associated with phosphorylated MAP1b that renders microtubules prone to depolymerisation, leading to the recovery of circa 3x less MAP1b than in WT (ESM_7).

Upon comparison of Kv_1.2_ and pY_458_Kv_1.2_ partners in an *Lgi1^-/-^* background (Fig. 3b), only four common partners were present: CCT4 (a chaperone T-complex subunit involved in actin and tubulin folding), glyceraldehyde-3-phosphate dehydrogenase (Gapdh/G3P) that can modulate the organization and assembly of the cytoskeleton, the glycolytic enzyme L-lactate dehydrogenase B chain (LDBH) indicator of an oxidative stress and the filamentous actin cross-linking protein myristoylated alanine-rich C-kinase substrate (MARCKS), which is the most prominent cellular substrate for protein kinase C. This comparison again suggests that Kv_1.2_ downregulation in *Lgi1^-/-^* is related to perturbation of association with the actin cytoskeleton. The presence of CCT4 at equivalent emPAI levels for phosphorylated and non-phospholytated Kv_1.2_ suggests that the absence of LGI1 have similar destabilizing effects on both channel types increasing their clearance [59].

### Partners of pY_458_Kv_1.2_ not associated with Kv_1.2_

pY_458_Kv_1.2_ is associated with several new partners (ESM_7, ESM_8 and ESM_9) that were not present in the Kv_1.2_ proteome [26]. Analysis of these partners in GO terms database shows a diverse molecular interactions and subcellular localisation (ESM_10) suggesting that despite a considerable difference in their proteomes, Kv_1_ channels containing pY_458_Kv_1.2_ are as widely distributed throughout neuronal compartments as the non-pY_458_Kv_1.2_.

## Discussion

In this study, an affinity purified specific Kv_1.2_pY_458_ antibody allowed us to identify minor high molecular weight Kv_1.2_ entities that were not detected previously. Among them a hyperglycosylated and tyrosine-phosphorylated (pY_458_) form of 100 kDa which is upregulated in *Lgi1^-/-^*. We did not detect any posttranslational modification in the other Kv_1.2_ entities that could explain their electrophoretic profile and therefore, their apparent high molecular weight may be attributed to aggregation despite the denaturation being performed at relatively low temperature.

Mass spectrometry analysis of protein complexes immunoprecipitated by pY_458_Kv_1.2_ antibodies from both wild type and *Lgi1^-/-^* backgrounds shows a significantly different profile from those previously described for Kv_1.2_ [26] with antibodies recognizing the same but non-phosphorylated Kv_1.2_ peptide (ESM_1). In addition, since the proteome of pY_458_Kv_1.2_ in *Lgi1^-/-^* background is different from that of Kv_1.2_ in *Lgi1^-/-^*, the phosphorylation of Y_458_ may not be the key parameter in Kv_1.2_ downregulation in *Lgi1^-/-^* and whether phosphorylation of Kv_1.2_ Y_458_ is linked to glycosylation cannot be inferred from the current study. Previous work showed that Kv1 glycosylation affects Kv_1_ gating properties [60] and facilitates channel trafficking to the cell membrane and its stabilization [61]. While a decrease in Kv_1.2_ expression level can be directly linked to the reduced D-type current and the increase in neuronal excitability in *Lgi1^-/-^*, the increase in the intensity of the described highly glycosylated form of Kv_1.2_ in *Lgi1^-/-^* questions the functional role of these subunits and their participation in the potassium conductance.

Recently, we uncovered the changes in expression and distribution of Kv_1.2_ in *Lgi1^-/-^* mice compared to their WT littermates [26]. However, a direct comparison between the staining patterns obtained with the monoclonal Kv_1.2_ and the polyclonal pY_458_Kv_1.2_ antibodies must be interpreted with caution. Moreover, citrate-buffer heat-induced antigen retrieval was performed for Kv_1.2_ but could not be applied to pKv_1.2_, as the phospho-epitope is labile and presumably dephosphorylates under such conditions. Nevertheless, although the overall staining distribution obtained with both antibodies was comparable throughout the hippocampal formation, three main differences were observed between the anti-Kv_1.2_ and the anti-pY_458_Kv_1.2_ staining. First, anti-Kv_1.2_ selectively labels axonal initial segments (AIS) of both, pyramidal cells and parvalbumin interneurons in the hippocampal formation, whereas such a staining pattern was not detected with the pY_458_Kv_1.2_ antibody. Second, Kv_1.2_ immunostaining was homogeneous through CA3 hippocampal region but markedly heterogeneous in CA1 and the Dentate Gyrus (DG), where it labelled preferentially the Stratum Lacunosum Moleculare (SLM) and the Molecular Layer (ML), respectively. In contrast, pY_458_-Kv_1.2_ was rather faint in the hippocampal formation, but showed preferential association with the stratum lucidum (SL) of CA3 and, to a lesser extent, the ML of the DG (see ESM_4). Tyrosine phosphorylation followed by internalization of Kv_1.2_ channels has been proposed as a mechanism mediating LTP of intrinsic excitability in CA3 pyramidal neurons [62] [63]. As such, the increased immunoreactivity in the SL of CA3, which is at least partially from a non-presynaptic origin (see Fig. 2), could underlie this phenomenon. Lastly, the pY_458_-Kv_1.2_, shows a prominent association with myelinated fiber bundles (ESM_4), a feature absents in Kv_1.2_ immunostainings.

We and others have shown that the detection of Kv_1_ channels by immunofluorescence is strongly influenced by Kv_1_ subcellular distribution and the molecular environment of the epitopes [64] [26]. Thus, a modified molecular environment of pY_458_Kv_1.2_ could partially account for the reduced accessibility of the antibody in WT compared to *Lgi1^-/-^*. While the staining of Kv_1.2_ and pY_458_-Kv_1.2_ can provide complementary information, they cannot be quantitatively compared in a straightforward manner. Any observed differences in staining intensity or distribution should therefore be interpreted as reflecting differences in epitope availability rather than absolute changes in protein quantity.

In contrast to the previously described Kv_1_._2_ channels subunit composition where several Kvα are part of the channel and the Kvβ_2_/Kvα_1.2_ ratio is >4 in WT animals [26], pY_458_Kv_1.2_ associated subunit composition is rather limited to Kv_1_._1_ and Kvβ_2_ in WT and Kvβ_2_ does not associate with Kv_1_._2_ in *Lgi1*^-/-^. In addition, Kvβ_2_/Kvα_1.2_ ratio is around 0.4 (ESM_1). The reason for the limited association of pY_458_Kv_1.2_ with other Kv_1_ subunits that are present in dendrotoxin sensitive channels is not clear. However, it may be speculated that this might be due to i) conformational changes induced by Y_458_ phosphorylation ii) the recruitment of new partners iii) changes in cytoskeletal links driving this modified subunit away from its partner subunits.

DLG2 and DLG4 were shown to associate with Kv_1.2_ [26] and PDZ interactions are crucial for Kv_1_ surface expression [17]. However, these interactions are dispensable for Kv_1.2_ juxtaparanodal clustering [65]. More recently a direct association of Kv_1.2_ with SCRIB was suggested to take place in the AIS and with PSD93 modulate AIS Kv1 clustering [27]. In central and peripheral myelinated neurons, the neuronal cell adhesion molecule Caspr2 and the neuro-glial glycosyl-phosphatidyl-inositol-anchored cell adhesion molecule TAG1 play an important role in juxtaparanodal clustering of Kv_1.2_ [66] [67] and homophilic cis and trans interactions of TAG1 are crucial in this mechanism [68], controlling thus the internodal resting potential [69] [66]. In this context, it was suggested that TAG1-mediated Kv_1.2_ clustering could be initiated by the enrichment of TAG1 in lipid rafts, followed by cis and trans homophilic TAG1 interactions and that this clustering is maintained by the local underlying actin cytoskeleton [68]. The phosphorylation of the C-terminal tyrosine residue Y_458_ in Kv_1.2_ is required for TAG1-mediated clustering mechanism [68] but reduces axonal targeting [17]. Therefore, one would expect pY_458_Kv_1.2_ to be less associated with axonal markers. Surprisingly, neither SCRIB nor Caspr2 or TAG1 are found in the list of pY_458_Kv_1.2_ partners and the newly identified channel population is not associated with any of the DLG proteins and is associated in WT with Kif5 and Kvβ_2_ suggesting their presence in an axonal pool [70] [71]. Altogether, the current data set suggests that the newly identified Kv_1.2_ population has fundamentally different molecular partnerships compared to the previously described one.

Immunofluorescence data corroborate the mass spectrometry findings and show that the hyperglycosylated pY_458_Kv_1.2_ occurs in axonal compartments. This localisation may be influenced by the hetero-multimeric nature of Kv_1_ channel subunits and their phosphorylation status. Kvβ is important in Kv_1.2_ trafficking [72] and the functional interaction of Kvα and Kvβ subunits depends on their respective phosphorylation status [73]. A continuum of S/T as well as Y residues phosphorylation may co-exist and therefore co-influence membrane expression, compartment targeting as well as binding to molecular partners.

Independently of Kvβ catalytic keto-reductase activity, its NADP^+^ binding property is important in Kv_1.2_ trafficking [72]. Only in *Lgi1^-/-^*, pY_458_Kv_1.2_ is associated with thioredoxin (emPAI 0.68) and aldolase (emPAI 0.43) (ESM_9), two metabolic enzymes linked to NADP. Interestingly, it was shown that aldolase may be inactivated by oxidative conditions and NADPH is able to reactivate it through the NADP-dependent thioredoxin system [74]. Together, the important recruitment of aldolase and thioredoxin to the proteome of pY_458_Kv_1.2_ may highlight a modification of oxidative stress in *Lgi1^-/-^* and suggest the existence of a specific metabolic pathway for pY_458_Kv_1.2_ in *Lgi1^-/-^*.

It is interesting to note that adducin that mediates actin-spectrin interaction [75] is found associated with pY_458_Kv_1.2_ only in *Lgi1^-/-^*. The phosphorylation of adducin disrupts adducin-mediated actin-spectrin interaction and leads to cytoskeletal reorganisation [76]. Knowing that actin territories control protrusions, while spectrin ones concentrate in retractile zones [77], this molecular interaction may be implicated in the cytoskeletal reorganisation in *Lgi1^-/-^* and the decrease in the number of dendritic spines.

Septins are GTP-binding cytoskeletal components that assemble to form hetero-oligomeric complexes, filaments, bundles and rings [78] [79]. They modulate the functions of actin and microtubules and act as scaffolds [79] for protein recruitment at the plasma membrane as well as in the cytosol. They bind to phosphoinositides and were shown to form diffusion barriers in dendritic spines [79] [80]. They also stabilize and maintain ankyrin G in the axonal initial segments [81]. In contrast to the association of Kv_1.2_ with septins in WT and *Lgi1^-/-^* as well as pY_458_Kv_1.2_ in WT, our current analysis shows the total loss of association pY_458_Kv_1.2_ with septins in *Lgi1^-/-^*. These data suggest that septins may play a crucial role in shaping the localisation of pY_458_Kv_1.2_.

Among the specific partners of pY_458_Kv_1.2_ that were absent from the Kv_1.2_ proteome, we found 2 proteins associated with the Rho GTPase effectors Arhgef2 and Arhgap26 in WT. In *Lgi1^-/-^*, the association with Arhgef2 is lost with a concomitant and strong association with 2 Rho proteins Rac1 and Rac3 [82] which is in agreement with the previously described significant increase in RhoA activity in *Lgi1^-/-^*. The Rho family members are molecular switches and their GTPase activity is crucial for an important number of cellular mechanisms from signal transduction to cytoskeleton remodelling [82] [83] [84]. The interaction of LGI1 with NgR1 reduces RhoA signalling promoting synapse formation and stabilization [11]. Also, RhoA plays a critical role in regulating the membrane flow through the endosomal system modulating the stability of the myelin proteolipid protein at oligodendroglial-membranes in contact with neurons [84]. Interestingly, GTP-bound RhoA was shown to co-imunoprecipitate with Kv_1.2_ and to regulate its activity since it plays a role in GPCR and tyrosine kinase-mediated downregulation of Kv_1.2_ activity [85].

The enzyme SIRT2 (NAD-dependent protein deacetylase sirtuin-2) is functionally associated with the tubulin cytoskeletal reorganisation through its implication in protein deacetylation [86]. The presence of SIRT2 in the proteome of only WT pY_458_Kv_1.2_ (ESM_8) may indicate an active deacetylation process that could affect the pY_458_Kv_1.2_ partner Tau (Fig. 1). Therefore, deacetylated Tau may have a reduced capacity to stabilize microtubules [87] leading to domains with impaired cytoskeletal stability where pY_458_Kv_1.2_ is enriched. Some common markers for the regulation of cytoskeletal dynamics and endocytic pathways were shared between Kv_1.2_ and pY_458_Kv_1.2_. While Cyfip2 was found specifically associated with Kv_1.2_ in WT (emPAI 0.11), it is a partner of pY_458_Kv_1.2_ in both WT and *Lgi1^-/-^* genotypes with comparable emPAI values (ESM_7) and this despite that the emPAI value of Kv_1.2_ immunoprecipitated by anti-pY_458_Kv_1.2_ is 10 times less than that immunoprecipitated by anti-Kv_1.2_ [26]. This may suggest that Cyfip2 is present in immunoprecipitated complexes through an interaction with a common partner to Kv_1.2_ and pY_458_Kv_1.2_ namely either Gsk3b or Ap1b1 (Fig. 3a, c and ESM_5). While Ap1b1 is an adaptor protein for endocytosis, Gsk3b is involved in cytoskeletal dynamics [88] and Cyfip2 is part of the WAVE1 complex and is involved in cytoskeletal dynamics and endocytic trafficking. AP2a2, and AP2b1, implicated in clathrin-mediated endocytosis, were equally enriched in both genotypes. AP3b2 and AP3d1 participate in polarized sorting [44] and their association with pY_458_Kv_1.2_ may indicate a common specific sorting pathway of the phosphorylated Kv_1.2_ subunit in both WT and *Lgi1^-/-^*. AP2m1 is responsible for cargo selection and directly interacts with ATP6V1H subunit and the autophagosome marker Map1lc3a [89] that are both present in the pY_458_Kv_1.2_ proteome from both genotypes. This may suggest that autophagosomes are part of the dynamic regulation of pY_458_Kv_1.2_ expression Annexin A2, copine 6 and 7 are calcium and lipid binding proteins involved in intracellular signal transduction [90] [91]. Also, annexin A2 is involved in endosomal repair following organelle destabilization [92] and copine 6 and 7 are SNARE binding proteins that monitor spontaneous neurotransmission [93] [94]. In *Lgi1^-/-^*, Kv_1.2_ is associated with copine 6 and annexin A2 [26] while pY_458_Kv_1.2_ only associates with the brain enriched copine 7. These data highlight again the different fate of pY_458_Kv_1.2_ between WT and *Lgi1^-/-^*.

We previously showed that Kv_1.2_ loses association with myelin associated proteins in *Lgi1^-/-^* [26] and suggested that one consequence of LGI1 depletion could be the exclusion of Kv_1.2_ from myelin containing regions. The association of pY_458_Kv_1.2_ with contactin 1, in both genotypes and PDIA3 and SIRT2 in WT may be an indicator of the presence of the newly identified form of Kv_1.2_ in myelinated axons [95]. However, the exclusive association of pY_458_Kv_1.2_ in *Lgi1^-/-^* with the myelin protein Cnp [96] as well as with SERA and ALDOA, that are present in the CNS myelin proteome [86], may participate stabilizing pY_458_Kv_1.2_ in the membrane of *Lgi1^-/-^*.

As for Kv_1.2,_ in *Lgi1^-/-^* the interactome of pY_458_Kv_1.2_ shows a major reshaping of its molecular neighbourhood. The major changes in molecules implicated in signalling cascade, the modifications in the association with proteins involved in trafficking and recycling dynamics, the increased links with metabolic/stress-related molecules argue that LGI1 behaves as a local organizer and its deletion results in the rewiring of intracellular Kv_1.2_ interactomes. The change in the association with the actin cytoskeleton in *Lgi1^-/-^* suggests a cytoskeletal re-anchoring /stabilization of Kv_1.2_ and pY_458_Kv_1.2_.

In conclusion, Since glycosylation facilitates Kv_1_ activation with lower depolarization intensities and stabilizes the channels in membranes [60] [61], the upregulated hyperglycosylated pY_458_Kv_1.2_ expression may be a homeostatic attempt from neurons to stabilize membrane Kv_1.2_ in response to LGI1 deletion. Of note, protein tyrosine phosphatase receptor type D (PTPRD) was found in a Genome-Wide Association study with LGI1-antibody encephalitis [97]. Therefore, one may speculate that inhibition of phospho-tyrosine kinase activity could be a way of stabilizing membrane Kv_1.2_ avoiding thus massive neuronal excitability perturbations. Future investigations should consider whether this could be a common attempt from biological systems to alleviate excitability increase due to Kv_1.2_ expression decrease or blockade.

## Acknowledgements

We are grateful for the Institut National de la Santé et de la Recherche Médicale INSERM), Aix-Marseille Université (AMU) and the Agence Nationale de la Recherche (ANR) for their financial support. We thank Stephanie Baulac for sharing the *Lgi1^−/−^* mice strain [98].

## Statements & declarations

### Author contributions

O.EF conceived the study, supervised the entire project, performed the experimental design, data analysis, classification, interpretation and manuscript preparation. M.Se. and C.L. contributed to the design of the study. J.R-F performed mass spectrometry data analysis, immunofluorescence experiments on brain slices, analysed images and prepared immunofluorescence figures. K.D. performed immunoprecipitations. J.R-F and K.D. took care of mice breeding and availability of *Lgi1^-/-^* animals. M. B. performed mass spectrometry experiments. O.EF and M.S. performed biochemical experiments. O.EF wrote the original draft of the manuscript and prepared the figures. All authors edited and reviewed the manuscript.

### Ethics approval

Not applicable

### Consent to participate

All authors approved submission

### Consent to publish

All authors approved publication

### Data and material availability

All data needed to evaluate the conclusions in the paper are present in the paper and/or the Supplementary Materials. The proteomic data were deposited on the PRIDE website (https://www.ebi.ac.uk/pride/archive) with the dataset identifier PXD 069032.

## References

1. Seagar M, Russier M, Caillard O, Maulet Y, Fronzaroli-Molinieres L, De San Feliciano M, et al. LGI1 tunes intrinsic excitability by regulating the density of axonal Kv1 channels. Proc Natl Acad Sci U S A. 2017;114:7719–24.

2. Extremet J, El Far O, Ankri N, Irani SR, Debanne D, Russier M. An Epitope-Specific LGI1-Autoantibody Enhances Neuronal Excitability by Modulating Kv1.1 Channel. Cells. 2022;11:

3. Petit-Pedrol M, Sell J, Planaguma J, Mannara F, Radosevic M, Haselmann H, et al. LGI1 antibodies alter Kv1.1 and AMPA receptors changing synaptic excitability, plasticity and memory. Brain : a journal of neurology. 2018;

4. Zhou L, Su LD, Cao SL, Xie YJ, Wang N, Shao CY, et al. Celecoxib Ameliorates Seizure Susceptibility in Autosomal Dominant Lateral Temporal Epilepsy. J Neurosci. 2018;38:3346–57.

5. Zhou L, Wang K, Xu Y, Dong BB, Wu DC, Wang ZX, et al. A patient-derived mutation of epilepsy-linked LGI1 increases seizure susceptibility through regulating K(v)1.1. Cell Biosci. 2023;13:34.

6. Ogawa Y, Oses-Prieto J, Kim MY, Horresh I, Peles E, Burlingame AL, et al. ADAM22, a Kv1 channel-interacting protein, recruits membrane-associated guanylate kinases to juxtaparanodes of myelinated axons. J Neurosci. 2010;30:1038–48.

7. Schulte U, Thumfart JO, Klocker N, Sailer CA, Bildl W, Biniossek M, et al. The epilepsy-linked Lgi1 protein assembles into presynaptic Kv1 channels and inhibits inactivation by Kvbeta1. Neuron. 2006;49:697–706.

8. Irani SR, Alexander S, Waters P, Kleopa KA, Pettingill P, Zuliani L, et al. Antibodies to Kv1 potassium channel-complex proteins leucine-rich, glioma inactivated 1 protein and contactin-associated protein-2 in limbic encephalitis, Morvan’s syndrome and acquired neuromyotonia. Brain : a journal of neurology. 2010;133:2734–48.

9. Ramirez-Franco J, Debreux K, Extremet J, Maulet Y, Belghazi M, Villard C, et al. Patient-derived antibodies reveal the subcellular distribution and heterogeneous interactome of LGI1. Brain : a journal of neurology. 2022;

10. Schiller MR. Coupling receptor tyrosine kinases to Rho GTPases--GEFs what’s the link. Cell Signal. 2006;18:1834–43.

11. Thomas RA, Gibon J, Chen CXQ, Chierzi S, Soubannier VG, Baulac S, et al. The Nogo Receptor Ligand LGI1 Regulates Synapse Number and Synaptic Activity in Hippocampal and Cortical Neurons. eNeuro. 2018;5:

12. Johnston J, Forsythe ID, Kopp-Scheinpflug C. Going native: voltage-gated potassium channels controlling neuronal excitability. J Physiol. 2010;588:3187–200.

13. Jan LY, Jan YN. Voltage-gated potassium channels and the diversity of electrical signalling. J Physiol. 2012;590:2591–9.

14. Debanne D, Campanac E, Bialowas A, Carlier E, Alcaraz G. Axon physiology. Physiol Rev. 2011;91:555–602.

15. Trimmer JS. Subcellular localization of K+ channels in mammalian brain neurons: remarkable precision in the midst of extraordinary complexity. Neuron. 2015;85:238–56.

16. Ogawa Y, Horresh I, Trimmer JS, Bredt DS, Peles E, Rasband MN. Postsynaptic density-93 clusters Kv1 channels at axon initial segments independently of Caspr2. J Neurosci. 2008;28:5731–9.

17. Gu C, Jan YN, Jan LY. A conserved domain in axonal targeting of Kv1 (Shaker) voltage-gated potassium channels. Science. 2003;301:646–9.

18. Inda MC, DeFelipe J, Munoz A. Voltage-gated ion channels in the axon initial segment of human cortical pyramidal cells and their relationship with chandelier cells. Proc Natl Acad Sci U S A. 2006;103:2920–5.

19. Browne DL, Gancher ST, Nutt JG, Brunt ER, Smith EA, Kramer P, et al. Episodic ataxia/myokymia syndrome is associated with point mutations in the human potassium channel gene, KCNA1. Nat Genet. 1994;8:136–40.

20. Herson PS, Virk M, Rustay NR, Bond CT, Crabbe JC, Adelman JP, et al. A mouse model of episodic ataxia type-1. Nat Neurosci. 2003;6:378–83.

21. Verdura E, Fons C, Schluter A, Ruiz M, Fourcade S, Casasnovas C, et al. Complete loss of KCNA1 activity causes neonatal epileptic encephalopathy and dyskinesia. J Med Genet. 2020;57:132–7.

22. Doring JH, Schroter J, Jungling J, Biskup S, Klotz KA, Bast T, et al. Refining Genotypes and Phenotypes in KCNA2-Related Neurological Disorders. Int J Mol Sci. 2021;22:

23. Hattan D, Nesti E, Cachero TG, Morielli AD. Tyrosine phosphorylation of Kv1.2 modulates its interaction with the actin-binding protein cortactin. J Biol Chem. 2002;277:38596–606.

24. Rasband MN, Park EW, Zhen D, Arbuckle MI, Poliak S, Peles E, et al. Clustering of neuronal potassium channels is independent of their interaction with PSD-95. J Cell Biol. 2002;159:663–72.

25. Duflocq A, Chareyre F, Giovannini M, Couraud F, Davenne M. Characterization of the axon initial segment (AIS) of motor neurons and identification of a para-AIS and a juxtapara-AIS, organized by protein 4.1B. BMC Biol. 2011;9:66.

26. Ramirez-Franco J, Debreux K, Sangiardi M, Belghazi M, Kim Y, Lee SH, et al. The downregulation of Kv(1) channels in Lgi1(-/-)mice is accompanied by a profound modification of its interactome and a parallel decrease in Kv(2) channels. Neurobiol Dis. 2024;196:106513.

27. Zhang W, Palfini VL, Wu Y, Ding X, Melton AJ, Gao Y, et al. A hierarchy of PDZ domain scaffolding proteins clusters the Kv1 K(+) channel protein complex at the axon initial segment. Sci Adv. 2025;11:eadv1281.

28. Nesti E, Everill B, Morielli AD. Endocytosis as a mechanism for tyrosine kinase-dependent suppression of a voltage-gated potassium channel. Mol Biol Cell. 2004;15:4073–88.

29. Perez-Riverol Y, Bandla C, Kundu DJ, Kamatchinathan S, Bai J, Hewapathirana S, et al. The PRIDE database at 20 years: 2025 update. Nucleic Acids Res. 2025;53:D543–D53.

30. Bolte S, Cordelieres FP. A guided tour into subcellular colocalization analysis in light microscopy. J Microsc. 2006;224:213–32.

31. Martinez-Marmol R, Styrczewska K, Perez-Verdaguer M, Vallejo-Gracia A, Comes N, Sorkin A, et al. Ubiquitination mediates Kv1.3 endocytosis as a mechanism for protein kinase C-dependent modulation. Sci Rep. 2017;7:42395.

32. Utsunomiya I, Tanabe S, Terashi T, Ikeno S, Miyatake T, Hoshi K, et al. Identification of amino acids in the pore region of Kv1.2 potassium channel that regulate its glycosylation and cell surface expression. J Neurochem. 2010;112:913–23.

33. Lorenzo DN, Edwards RJ, Slavutsky AL. Spectrins: molecular organizers and targets of neurological disorders. Nat Rev Neurosci. 2023;24:195–212.

34. Stirling L, Williams MR, Morielli AD. Dual roles for RHOA/RHO-kinase in the regulated trafficking of a voltage-sensitive potassium channel. Mol Biol Cell. 2009;20:2991–3002.

35. Prosser DC, Tran D, Gougeon PY, Verly C, Ngsee JK. FFAT rescues VAPA-mediated inhibition of ER-to-Golgi transport and VAPB-mediated ER aggregation. J Cell Sci. 2008;121:3052–61.

36. Sandhu J, Li S, Fairall L, Pfisterer SG, Gurnett JE, Xiao X, et al. Aster Proteins Facilitate Nonvesicular Plasma Membrane to ER Cholesterol Transport in Mammalian Cells. Cell. 2018;175:514–29 e20.

37. James C, Kehlenbach RH. The Interactome of the VAP Family of Proteins: An Overview. Cells. 2021;10:

38. Robinson RC, Turbedsky K, Kaiser DA, Marchand JB, Higgs HN, Choe S, et al. Crystal structure of Arp2/3 complex. Science. 2001;294:1679–84.

39. Lee M, Liu YC, Chen C, Lu CH, Lu ST, Huang TN, et al. Ecm29-mediated proteasomal distribution modulates excitatory GABA responses in the developing brain. J Cell Biol. 2020;219:

40. Galic M, Jeong S, Tsai FC, Joubert LM, Wu YI, Hahn KM, et al. External push and internal pull forces recruit curvature-sensing N-BAR domain proteins to the plasma membrane. Nat Cell Biol. 2012;14:874–81.

41. Gilligan DM, Lozovatsky L, Gwynn B, Brugnara C, Mohandas N, Peters LL. Targeted disruption of the beta adducin gene (Add2) causes red blood cell spherocytosis in mice. Proc Natl Acad Sci U S A. 1999;96:10717–22.

42. She BR, Liou GG, Lin-Chao S. Association of the growth-arrest-specific protein Gas7 with F-actin induces reorganization of microfilaments and promotes membrane outgrowth. Exp Cell Res. 2002;273:34–44.

43. Spiliotis ET, Nakos K. Cellular functions of actin- and microtubule-associated septins. Curr Biol. 2021;31:R651–R66.

44. Blumstein J, Faundez V, Nakatsu F, Saito T, Ohno H, Kelly RB. The neuronal form of adaptor protein-3 is required for synaptic vesicle formation from endosomes. J Neurosci. 2001;21:8034–42.

45. van den Boom J, Meyer H. VCP/p97-Mediated Unfolding as a Principle in Protein Homeostasis and Signaling. Mol Cell. 2018;69:182–94.

46. Bos JL, Rehmann H, Wittinghofer A. GEFs and GAPs: critical elements in the control of small G proteins. Cell. 2007;129:865–77.

47. Armstrong MC, Weiss YR, Hoachlander-Hobby LE, Roy AA, Visco I, Moe A, et al. The biochemical mechanism of Rho GTPase membrane binding, activation and retention in activity patterning. Embo J. 2025;44:2620–57.

48. de Curtis I. The Rac3 GTPase in Neuronal Development, Neurodevelopmental Disorders, and Cancer. Cells. 2019;8:

49. Tyckaert F, Zanin N, Morsomme P, Renard HF. Rac1, the actin cytoskeleton and microtubules are key players in clathrin-independent endophilin-A3-mediated endocytosis. J Cell Sci. 2022;135:

50. Adams DR, Ron D, Kiely PA. RACK1, A multifaceted scaffolding protein: Structure and function. Cell Commun Signal. 2011;9:22.

51. Myllykoski M, Seidel L, Muruganandam G, Raasakka A, Torda AE, Kursula P. Structural and functional evolution of 2’,3’-cyclic nucleotide 3’-phosphodiesterase. Brain Res. 2016;1641:64–78.

52. Rosener M, Muraro PA, Riethmuller A, Kalbus M, Sappler G, Thompson RJ, et al. 2’,3’-cyclic nucleotide 3’-phosphodiesterase: a novel candidate autoantigen in demyelinating diseases. J Neuroimmunol. 1997;75:28–34.

53. Aston C, Jiang L, Sokolov BP. Transcriptional profiling reveals evidence for signaling and oligodendroglial abnormalities in the temporal cortex from patients with major depressive disorder. Mol Psychiatry. 2005;10:309–22.

54. Sequeira A, Mamdani F, Ernst C, Vawter MP, Bunney WE, Lebel V, et al. Global brain gene expression analysis links glutamatergic and GABAergic alterations to suicide and major depression. PLoS One. 2009;4:e6585.

55. Peirce TR, Bray NJ, Williams NM, Norton N, Moskvina V, Preece A, et al. Convergent evidence for 2’,3’-cyclic nucleotide 3’-phosphodiesterase as a possible susceptibility gene for schizophrenia. Arch Gen Psychiatry. 2006;63:18–24.

56. Che R, Tang W, Zhang J, Wei Z, Zhang Z, Huang K, et al. No relationship between 2’,3’-cyclic nucleotide 3’-phosphodiesterase and schizophrenia in the Chinese Han population: an expression study and meta-analysis. BMC Med Genet. 2009;10:31.

57. Goold RG, Owen R, Gordon-Weeks PR. Glycogen synthase kinase 3beta phosphorylation of microtubule-associated protein 1B regulates the stability of microtubules in growth cones. J Cell Sci. 1999;112 ( Pt 19):3373–84.

58. Halpain S, Dehmelt L. The MAP1 family of microtubule-associated proteins. Genome Biol. 2006;7:224.

59. Ma X, Lu C, Chen Y, Li S, Ma N, Tao X, et al. CCT2 is an aggrephagy receptor for clearance of solid protein aggregates. Cell. 2022;185:1325–45 e22.

60. Watanabe I, Zhu J, Sutachan JJ, Gottschalk A, Recio-Pinto E, Thornhill WB. The glycosylation state of Kv1.2 potassium channels affects trafficking, gating, and simulated action potentials. Brain Res. 2007;1144:1–18.

61. Thayer DA, Yang SB, Jan YN, Jan LY. N-linked glycosylation of Kv1.2 voltage-gated potassium channel facilitates cell surface expression and enhances the stability of internalized channels. J Physiol. 2016;594:6701–13.

62. Hyun JH, Eom K, Lee KH, Ho WK, Lee SH. Activity-dependent downregulation of D-type K+ channel subunit Kv1.2 in rat hippocampal CA3 pyramidal neurons. J Physiol. 2013;591:5525–40.

63. Eom K, Kim Y, Lee SH. Input-specific bidirectional regulation of hippocampal CA3 pyramidal cell excitability. J Physiol. 2025;603:4005–25.

64. Lorincz A, Nusser Z. Specificity of immunoreactions: the importance of testing specificity in each method. J Neurosci. 2008;28:9083–6.

65. Horresh I, Poliak S, Grant S, Bredt D, Rasband MN, Peles E. Multiple molecular interactions determine the clustering of Caspr2 and Kv1 channels in myelinated axons. J Neurosci. 2008;28:14213–22.

66. Traka M, Goutebroze L, Denisenko N, Bessa M, Nifli A, Havaki S, et al. Association of TAG-1 with Caspr2 is essential for the molecular organization of juxtaparanodal regions of myelinated fibers. J Cell Biol. 2003;162:1161–72.

67. Pinatel D, Hivert B, Saint-Martin M, Noraz N, Savvaki M, Karagogeos D, et al. The Kv1-associated molecules TAG-1 and Caspr2 are selectively targeted to the axon initial segment in hippocampal neurons. J Cell Sci. 2017;130:2209–20.

68. Gu C, Gu Y. Clustering and activity tuning of Kv1 channels in myelinated hippocampal axons. J Biol Chem. 2011;286:25835–47.

69. Poliak S, Salomon D, Elhanany H, Sabanay H, Kiernan B, Pevny L, et al. Juxtaparanodal clustering of Shaker-like K+ channels in myelinated axons depends on Caspr2 and TAG-1. J Cell Biol. 2003;162:1149–60.

70. Rivera J, Chu PJ, Lewis TL, Jr., Arnold DB. The role of Kif5B in axonal localization of Kv1 K(+) channels. Eur J Neurosci. 2007;25:136–46.

71. Pinatel D, Faivre-Sarrailh C. Assembly and Function of the Juxtaparanodal Kv1 Complex in Health and Disease. Life (Basel). 2020;11:

72. Campomanes CR, Carroll KI, Manganas LN, Hershberger ME, Gong B, Antonucci DE, et al. Kv beta subunit oxidoreductase activity and Kv1 potassium channel trafficking. J Biol Chem. 2002;277:8298–305.

73. Navarro-Perez M, Estadella I, Benavente-Garcia A, Orellana-Fernandez R, Petit A, Ferreres JC, et al. The Phosphorylation of Kv1.3: A Modulatory Mechanism for a Multifunctional Ion Channel. Cancers (Basel). 2023;15:

74. van der Linde K, Gutsche N, Leffers HM, Lindermayr C, Muller B, Holtgrefe S, et al. Regulation of plant cytosolic aldolase functions by redox-modifications. Plant Physiol Biochem. 2011;49:946–57.

75. Fowler VM. The human erythrocyte plasma membrane: a Rosetta Stone for decoding membrane-cytoskeleton structure. Curr Top Membr. 2013;72:39–88.

76. Bennett V, Baines AJ. Spectrin and ankyrin-based pathways: metazoan inventions for integrating cells into tissues. Physiol Rev. 2001;81:1353–92.

77. Ghisleni A, Galli C, Monzo P, Ascione F, Fardin MA, Scita G, et al. Complementary mesoscale dynamics of spectrin and acto-myosin shape membrane territories during mechanoresponse. Nat Commun. 2020;11:5108.

78. Radler MR, Spiliotis ET. Right place, right time - Spatial guidance of neuronal morphogenesis by septin GTPases. Curr Opin Neurobiol. 2022;75:102557.

79. Benoit B, Pous C, Baillet A. Septins as membrane influencers: direct play or in association with other cytoskeleton partners. Front Cell Dev Biol. 2023;11:1112319.

80. Mostowy S, Cossart P. Septins: the fourth component of the cytoskeleton. Nat Rev Mol Cell Biol. 2012;13:183–94.

81. Hamdan H, Lim BC, Torii T, Joshi A, Konning M, Smith C, et al. Mapping axon initial segment structure and function by multiplexed proximity biotinylation. Nat Commun. 2020;11:100.

82. Mosaddeghzadeh N, Ahmadian MR. The RHO Family GTPases: Mechanisms of Regulation and Signaling. Cells. 2021;10:

83. Hossen E, Funahashi Y, Faruk MO, Ahammad RU, Amano M, Yamada K, et al. Rho-Kinase/ROCK Phosphorylates PSD-93 Downstream of NMDARs to Orchestrate Synaptic Plasticity. Int J Mol Sci. 2022;24:

84. Kippert A, Trajkovic K, Rajendran L, Ries J, Simons M. Rho regulates membrane transport in the endocytic pathway to control plasma membrane specialization in oligodendroglial cells. J Neurosci. 2007;27:3560–70.

85. Cachero TG, Morielli AD, Peralta EG. The small GTP-binding protein RhoA regulates a delayed rectifier potassium channel. Cell. 1998;93:1077–85.

86. Werner HB, Kuhlmann K, Shen S, Uecker M, Schardt A, Dimova K, et al. Proteolipid protein is required for transport of sirtuin 2 into CNS myelin. J Neurosci. 2007;27:7717–30.

87. Cook C, Stankowski JN, Carlomagno Y, Stetler C, Petrucelli L. Acetylation: a new key to unlock tau’s role in neurodegeneration. Alzheimers Res Ther. 2014;6:29.

88. Hajka D, Budziak B, Pietras L, Duda P, McCubrey JA, Gizak A. GSK3 as a Regulator of Cytoskeleton Architecture: Consequences for Health and Disease. Cells. 2021;10:

89. Tian Y, Chang JC, Fan EY, Flajolet M, Greengard P. Adaptor complex AP2/PICALM, through interaction with LC3, targets Alzheimer’s APP-CTF for terminal degradation via autophagy. Proc Natl Acad Sci U S A. 2013;110:17071–6.

90. Tang H, Pang P, Qin Z, Zhao Z, Wu Q, Song S, et al. The CPNE Family and Their Role in Cancers. Front Genet. 2021;12:689097.

91. Khvotchev M, Soloviev M. Copines, a Family of Calcium Sensor Proteins and Their Role in Brain Function. Biomolecules. 2024;14:

92. Scharf B, Clement CC, Wu XX, Morozova K, Zanolini D, Follenzi A, et al. Annexin A2 binds to endosomes following organelle destabilization by particulate wear debris. Nat Commun. 2012;3:755.

93. Liu P, Khvotchev M, Li YC, Chanaday NL, Kavalali ET. Copine-6 Binds to SNAREs and Selectively Suppresses Spontaneous Neurotransmission. J Neurosci. 2018;38:5888–99.

94. Savino M, d’Apolito M, Centra M, van Beerendonk HM, Cleton-Jansen AM, Whitmore SA, et al. Characterization of copine VII, a new member of the copine family, and its exclusion as a candidate in sporadic breast cancers with loss of heterozygosity at 16q24.3. Genomics. 1999;61:219–26.

95. Boyle ME, Berglund EO, Murai KK, Weber L, Peles E, Ranscht B. Contactin orchestrates assembly of the septate-like junctions at the paranode in myelinated peripheral nerve. Neuron. 2001;30:385–97.

96. Raasakka A, Kursula P. Flexible Players within the Sheaths: The Intrinsically Disordered Proteins of Myelin in Health and Disease. Cells. 2020;9:

97. Binks SNM, Elliott KS, Muniz-Castrillo S, Gilbert E, Kawasaki de Araujo T, Harper AR, et al. Novel risk loci in LGI1-antibody encephalitis: genome-wide association study discovery and validation cohorts. Brain : a journal of neurology. 2025;148:737–45.

98. Chabrol E, Navarro V, Provenzano G, Cohen I, Dinocourt C, Rivaud-Pechoux S, et al. Electroclinical characterization of epileptic seizures in leucine-rich, glioma-inactivated 1-deficient mice. Brain : a journal of neurology. 2010;133:2749–62.

