## Supplementary figures for "A hyperglycosylated form of Kv_1.2_ upregulated in LGI1 knockout mice"

**A**

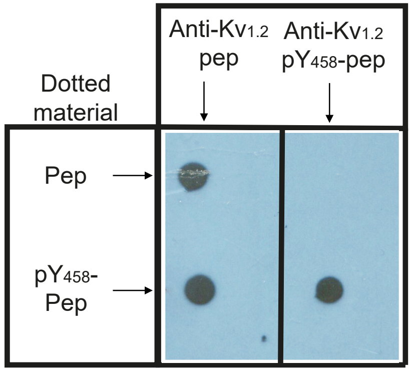


**ESM_1**

**Specificity of the $pY_{458}Kv_{1.2}$ antibody. a** Dot blot showing the specific recognition by the $pY_{458}Kv_{1.2}$ antibody of the phosphorylated immunogen compared to the non-phosphorylated peptide. **b** Alignment of the C-terminal sequences of mice $Kv_{1.2}$, $Kv_{1.1}$, $Kv_{1.3}$ $Kv_{1.4}$, $Kv_{1.5}$ and $Kv_{1.6}$. The immunogen peptide sequence is boxed in black and the epitope of the Neuromab K14/16 in a dashed box. The phosphorylated $Y_{458}$ residue is marked in red. **c** Comparison of $Kv_1$ subunits recovery by mass spectrometry analysis after immunoprecipitation from WT and *Lgi1*$^{-/-}$ using $pKv_{1.2}$ [26] and $pY_{458}Kv_{1.2}$ antibodies.

**B**

$pY_{458}Kv_{1.2}$ epitope (454-468)

Neuromab K14/16 epitope (467-484)

```
      411                                                                                       499
Kv1.2 SNFNYFYHRETEGEEQAQYLQ-VTSCPKIPSS-P-DLKKSRSASTISKSDY------MEIQEGVNNSNEDFREENLKTANCTL------ANTN-----YVNITKMLTDV
Kv1.1 SNFNYFYHRETEGEEQAQLLH-VSS-PNLASD-S-DLS-RRSSSTISKSEY------MEIEEDMNNSIAHYRQANIRTGNCTA------TDQN-----CVNKSKLLTDV
Kv1.3 SNFNYFYHRETEGEEQAQYMH-VGSCQHLSSSAE-ELRKARSNSTLSKSEY------MVIEEGGMNHS-AFPQTPFKTGNSTATCTTNNNPNS-----CVNIKKIFTDV
Kv1.4 SNFNYFYHRETENEEQTQLTQNAVSCPYLPSNLLKKFRSSTSSSLGDKSEY------LEMEEGVKESLCGKEEKCQGKGDD-----SETDKNN-----CSNAKAVETDV
Kv1.5 SNFNYFYHRETDHEEQAALKEEQGNQRR-ESGLDTGGQ---RKVSCSKASFCKTGGSLESSDSIRRGSCPLEKCHLKAK-------SNVDLRRSLYALCLDT-SRETDL
Kv1.6 SNFNYFYHRETEQEEQGQYTHVTCGQPT--PDL----K-A-TDNGLGKPDF------AEASRERRSSYLP------------------TPHR-----AYAEKRMLTEV
      ***********: ***     .   .      .           .* .:         .                                      *::
```

**C**

| Identification of $Kv_1$ isoforms associated with $Kv_{1.2}$ and $pY_{458}Kv_{1.2}$ in WT and *Lgi1* $^{-/-}$ | | | | |
|---|---|---|---|---|
| | $Kv_{1.2}$ WT | $pY_{458}Kv_{1.2}$ WT | $Kv_{1.2}$ *Lgi1*$^{-/-}$ | $pY_{458}Kv_{1.2}$ *Lgi1*$^{-/-}$ |
| **Kvα** | $Kv_{1.1}$; $Kv_{1.2}$; $Kv_{1.3}$ $Kv_{1.4}$; $Kv_{1.5}$; $Kv_{1.6}$ | $Kv_{1.1}$; $Kv_{1.2}$ | $Kv_{1.1}$; $Kv_{1.2}$; $Kv_{1.3}$ $Kv_{1.4}$; $Kv_{1.5}$; $Kv_{1.6}$ | $Kv_{1.2}$ |
| **emPAI** | 2.8; 2.9; 3.1; 1.5; 0.4; 3.3 | 0.26; 0.36 | 2.9; 2.8; 2.7; 1.3; 0.4; 2.2 | 0.14 |
| **Kvβ** | $Kv\beta_1$; $Kv\beta_2$ | $Kv\beta_2$ | $Kv\beta_1$; $Kv\beta_2$ | - |
| **emPAI** | 3.4; 14 | 0.16 | 2.9; 12 | - |

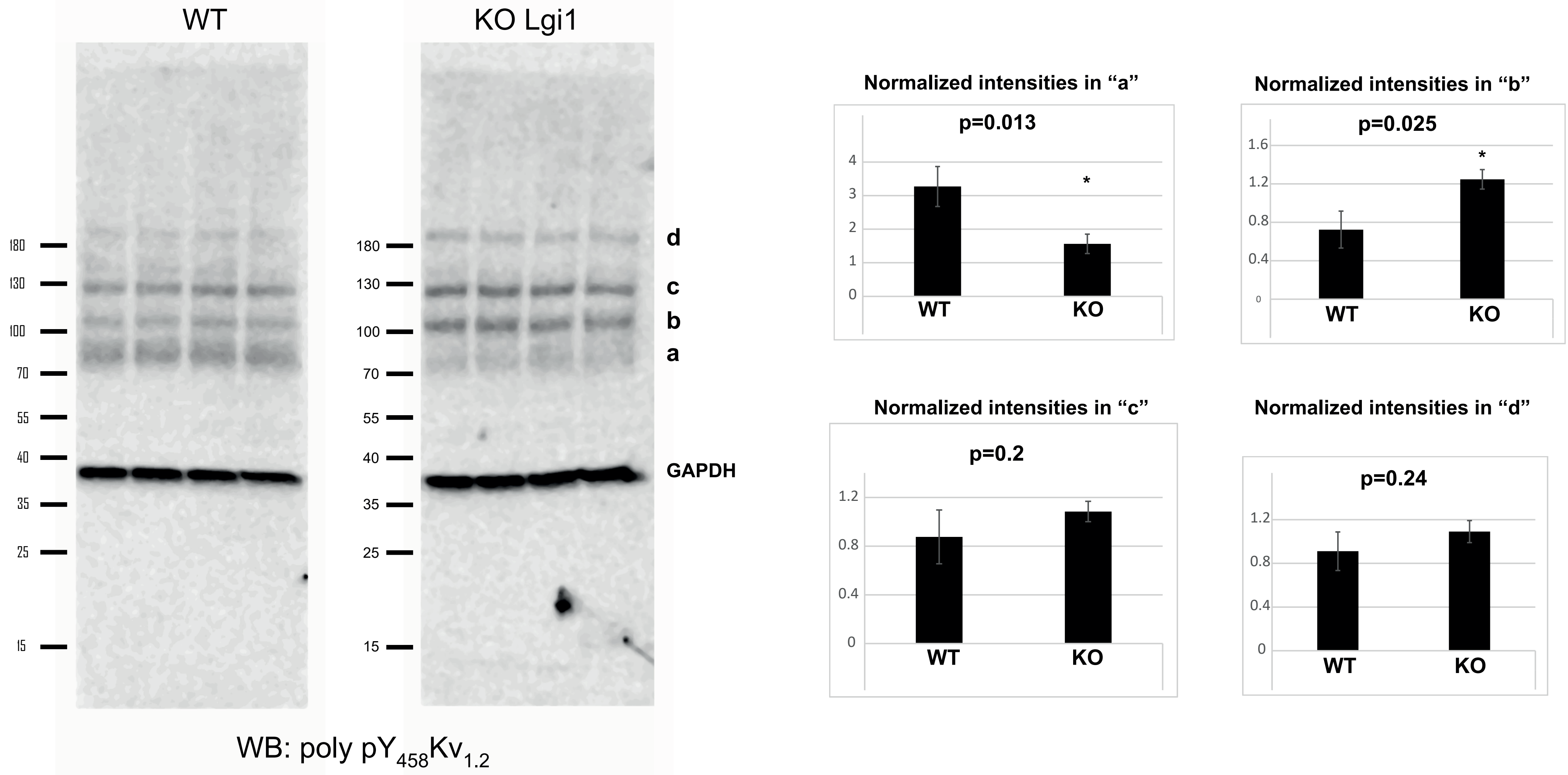


**ESM_2**

**Characterisation by Western blot of $pY_{458}Kv_{1.2}$ antibody a** Total mice brain homogenate from WT and *Lgi1*$^{-/-}$ were analysed by Western blot using $pY_{458}Kv_{1.2}$ antibody. **b** The four principal bands recognized by the antibody in (**a**) were quantified and their expression level difference between WT and *Lgi1*$^{-/-}$ were normalized to GAPDH and plotted in histograms. * indicate significant statistical differences

**ESM_3**

Post-translational modifications of $pY_{458}Kv_{1.2}$ immunoprecipitated material. Western blot with anti-ubiquitin and anti-SUMO2/3 antibodies after imununoprecipitation with $pY_{458}Kv_{1.2}$ from solubilized WT and *Lgi1*$^{-/-}$ brain extracts.

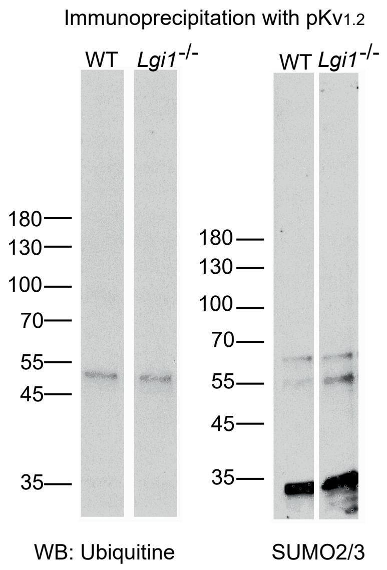

**ESM_4**

**a** Representative full-width coronal brain section from a WT mouse at Bregma ~–2.50 mm, immunostained for phosphorylated Kv$_{1.2}$ (pY$_{458}$Kv$_{1.2}$, green), AnkyrinG (red), and DAPI (blue), with merged images shown. Scale bar = 1 mm. **b** The same section highlighting pY$_{458}$Kv$_{1.2}$ (green) immunoreactivity within prominent myelinated fiber tracts. Notable neuroanatomical landmarks with intense labeling include the corpus callosum (cc), habenular commissure (hc), brachium of the superior colliculus (bsc), superior thalamic radiation (str), fimbria (fi), fasciculus retroflexus (fr), medial lemniscus (ml), cerebral peduncle (cp), mammillary tracts (mt), and the optic tract (opt). These structures exhibit robust signal consistent with pY$_{458}$Kv$_{1.2}$ localization along compact, highly myelinated axonal bundles. **c** High-magnification views of selected annotated regions shown in B, illustrating detailed pY$_{458}$Kv$_{1.2}$ distribution within each tract. Scale bar = 100 µm.

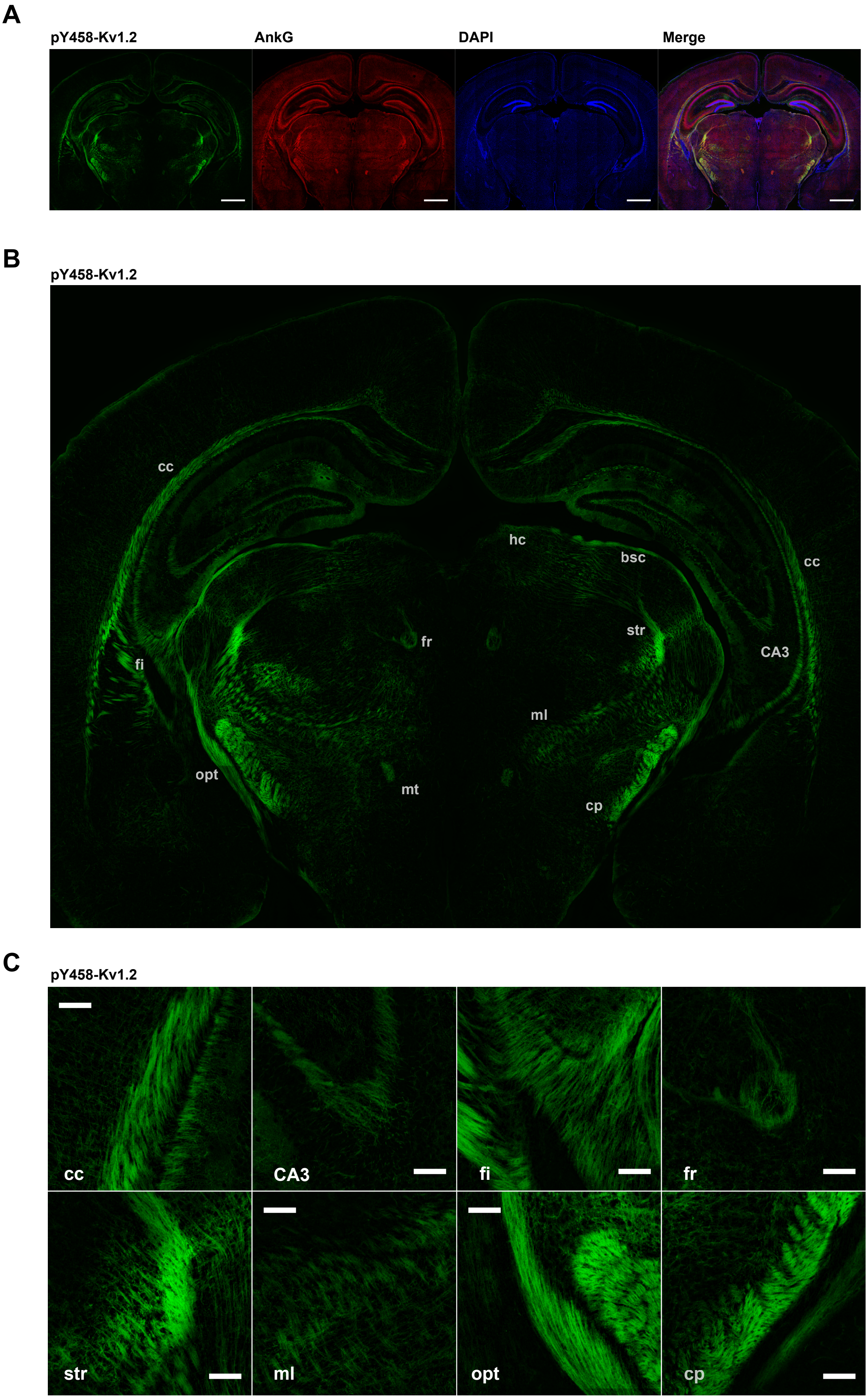

### ESM_5

#### Common partners to $Kv_{1.2}$ and $pY_{458}Kv_{1.2}$ and their Gene Ontology classification

| | Common partners between $Kv_{1.2}$ and $pY_{458}Kv_{1.2}$ |
|---|---|
| In WT | **Kcna2;Ap2a2;Atp1a1;Gramd1b;Kif2a;Slc1a2;Vapa;** Ap1b1;Gsk3b;Kcna1;Kcnab2;Ppp3ca; Ppp3cb; Ywhae; Cyfip2 |
| In *Lgi1*$^{-/-}$ | **Kcna2;Ap2a2;Atp1a1;Gramd1b;Kif2a;Slc1a2;Vapa;** Cct4;Dcx;G3P;Hsd17b4;Add1;Ldhb;Lancl2;Marcks;Stx1b |

### In WT

Cellular Component (Gene Ontology)

| GO-term | description | count in network | strength | false discovery rate |
|---|---|---|---|---|
| GO:0005955 | Calcineurin complex | 2 of 6 | 2.96 | 0.0017 |
| GO:0044224 | Juxtaparanode region of axon | 2 of 10 | 2.74 | 0.0026 |
| GO:0043194 | Axon initial segment | 2 of 27 | 2.31 | 0.0108 |
| GO:0008076 | Voltage-gated potassium channel complex | 2 of 76 | 1.86 | 0.0386 |
| GO:0098978 | Glutamatergic synapse | 5 of 522 | 1.42 | 0.00043 |
| GO:0150034 | Distal axon | 4 of 420 | 1.41 | 0.0027 |
| GO:0098794 | Postsynapse | 4 of 801 | 1.13 | 0.0166 |
| GO:0045202 | Synapse | 7 of 1589 | 1.08 | 0.00017 |
| GO:0099512 | Supramolecular fiber | 4 of 1064 | 1.01 | 0.0386 |
| GO:0043005 | Neuron projection | 6 of 1721 | 0.98 | 0.0024 |
| GO:0070161 | Anchoring junction | 4 of 1193 | 0.96 | 0.0468 |
| GO:0005829 | Cytosol | 7 of 4213 | 0.66 | 0.0126 |
| GO:0032991 | Protein-containing complex | 8 of 5743 | 0.58 | 0.0059 |

Subcellular localization (COMPARTMENTS)

| compartment | description | count in network | strength | false discovery rate |
|---|---|---|---|---|
| GOCC:0150034 | Distal axon | 4 of 354 | 1.49 | 0.0104 |
| GOCC:0099512 | Supramolecular fiber | 4 of 692 | 1.2 | 0.0142 |
| GOCC:0098794 | Postsynapse | 4 of 598 | 1.26 | 0.0130 |
| GOCC:0044224 | Juxtaparanode region of axon | 2 of 11 | 2.7 | 0.0104 |
| GOCC:0043194 | Axon initial segment | 2 of 26 | 2.32 | 0.0130 |
| GOCC:0043005 | Neuron projection | 5 of 1122 | 1.09 | 0.0104 |
| GOCC:0032991 | Protein-containing complex | 8 of 5305 | 0.61 | 0.0104 |
| GOCC:0008076 | Voltage-gated potassium channel complex | 2 of 70 | 1.89 | 0.0385 |
| GOCC:0005963 | Magnesium-dependent protein serine/threonine phosphata... | 2 of 28 | 2.29 | 0.0130 |
| GOCC:0005955 | Calcineurin complex | 2 of 8 | 2.83 | 0.0104 |
| GOCC:0005829 | Cytosol | 5 of 2014 | 0.83 | 0.0385 |

##### In *Lgi1*$^{-/-}$

Cellular Component (Gene Ontology)

| GO-term | description | count in network | strength | false discovery rate |
|---|---|---|---|---|
| GO:0005856 | Cytoskeleton | 7 of 2401 | 0.85 | 0.0118 |

Subcellular localization (COMPARTMENTS)

| compartment | description | count in network | strength | false discovery rate |
|---|---|---|---|---|
| GOCC:0099513 | Polymeric cytoskeletal fiber | 4 of 456 | 1.33 | 0.0248 |
| GOCC:0005856 | Cytoskeleton | 6 of 1564 | 0.97 | 0.0209 |
| GOCC:0005737 | Cytoplasm | 9 of 7282 | 0.48 | 0.0376 |

#### ESM_6

a. Exclusive and common partners of $Kv_{1.2}$ in WT or *Lgi1*$^{-/-}$

b. Exclusive and common partners of $pY_{458}Kv_{1.2}$ in WT or *Lgi1*$^{-/-}$

**A**

### Kv1.2 partners in WT versus LGI1 KO (emPAI>=0.1)

**Exclusive to WT**

| | | | |
|---|---|---|---|
| ACTC | 2.12 | DTX1 | 0.13 |
| AP3S1 | 0.26 | ATP2B4 | 0.12 |
| Plp1 (Mypr) | 0.23 | RANBP9 | 0.12 |
| Ppp3cb | 0.22 | MYH10 | 0.12 |
| AP1B1 | 0.20 | GSK3B | 0.12 |
| DLG2 | 0.15 | Gramd1a | 0.11 |
| LGI1 | 0.14 | cyfip2 | 0.11 |
| VIME | 0.13 | GSDMA | 0.11 |

**Exclusive to LGI1 KO**

| | | | |
|---|---|---|---|
| G3P | 1.64 | Syngr3 | 0.22 |
| FABP7 | 0.81 | LRRC59 | 0.21 |
| VAMP2 | 0.48 | STX1A | 0.21 |
| MARCKS | 0.41 | SYN2 | 0.21 |
| PCMTD2 | 0.32 | LDHB | 0.19 |
| VDAC1 | 0.31 | Prkacb | 0.16 |
| SEPTIN7 | 0.30 | RACK1 | 0.16 |
| CPNE6 | 0.29 | DNM3 | 0.15 |
| CCT4 | 0.27 | SEPTIN6 | 0.14 |
| Ythdf1 | 0.27 | Lancl2 | 0.10 |
| ANXA2 | 0.24 | RPN1 | 0.10 |
| DCX | 0.23 | | |

**Common partners with emPAI ratios WT/ LGI1 KO**

| | | | | | |
|---|---|---|---|---|---|
| ADAM22 | 2.85 | KCNAB1 | 1.19 | TRPV2 | 0.89 |
| DLG4 | 2.04 | KCNA4 | 1.19 | Atp1a1 | 0.88 |
| DLG1 | 1.80 | KCNA3 | 1.16 | Hsd17b4 | 0.85 |
| DLG3 | 1.67 | KCNAB2 | 1.16 | WDR26 | 0.82 |
| PSD | 1.63 | VAPB | 1.08 | SEPTIN11 | 0.80 |
| ATP2B2 | 1.60 | Epm2aip1 | 1.06 | STX1B | 0.78 |
| ADD1 | 1.58 | Csnk1e | 1.04 | Ppp3ca | 0.74 |
| KCNA6 | 1.49 | KCNA2 | 1.03 | LRTM2 | 0.73 |
| Prkcb | 1.44 | AP2A2 | 1.01 | Lap3 | 0.68 |
| PACS1 | 1.42 | KCNA5 | 0.99 | Ywhae | 0.64 |
| ATP1A2 | 1.25 | Gramd1b | 0.98 | VAPA | 0.56 |
| WNK2 | 1.21 | KCNA1 | 0.96 | Ywhag | 0.44 |
| Slc1a2 | 1.21 | LARGE1 | 0.96 | Ythdf1 | 0.31 |
| KIF2A | 1.20 | Aldh18a1 | 0.95 | SEPTIN8 | 0.14 |

**B**

### pY458Kv1.2 partners in WT versus LGI1 KO (emPAI>=0.1)

**Exclusive to WT**

| | | | | | |
|---|---|---|---|---|---|
| PIN1 | 1.23 | CDK5 | 0.29 | Septin5 | 0.16 |
| NEUG | 1.18 | SEC22B | 0.29 | BTBD17 | 0.15 |
| CYCS | 1.04 | PDIA3 | 0.28 | Rab11fip2 | 0.15 |
| GNAO | 0.90 | MRP | 0.28 | PGK1 | 0.15 |
| Ywhae | 0.78 | PFKM | 0.27 | Arhgef2 | 0.14 |
| TAU | 0.63 | KCNA1 | 0.27 | Septin11 | 0.14 |
| DNM1 | 0.60 | SNAP25 | 0.24 | Septin8 | 0.14 |
| DIRAS2 | 0.51 | RACK1 | 0.22 | WASF1 | 0.14 |
| DCLK1 | 0.46 | Acot7 | 0.21 | ABI1 | 0.14 |
| OTUB1 | 0.43 | LDHA | 0.21 | CORO1C | 0.13 |
| Ppp3ca | 0.41 | KCTD12 | 0.19 | Sptbn1 | 0.12 |
| GNB2 | 0.39 | SIRT2 | 0.19 | ASAP1 | 0.11 |
| BAIAP2 | 0.38 | VPS35 | 0.19 | Slc1a3 | 0.11 |
| CSNK2B | 0.35 | PON1 | 0.17 | PFKL | 0.10 |
| Sptan1 | 0.32 | Ppp2r1a | 0.17 | G3BP2 | 0.10 |
| CFL1 | 0.31 | GNB5 | 0.16 | | |
| MAPK3 | 0.30 | KCNAB2 | 0.16 | | |

**Exclusive to LGI1 KO**

| | | | | | |
|---|---|---|---|---|---|
| G3P | 1.53 | CRIP2 | 0.31 | Actr1a | 0.16 |
| MT3 | 1.03 | FHL1 | 0.29 | KAP3 | 0.16 |
| RAC1 | 0.93 | STX1B | 0.27 | PFKP | 0.16 |
| ATP1B1 | 0.90 | SYP | 0.27 | UBA1 | 0.15 |
| RAC3 | 0.82 | GAS7 | 0.25 | VCP | 0.14 |
| MARCKS | 0.72 | PRKCG | 0.25 | SEPTIN3 | 0.13 |
| TXN | 0.68 | CYRIB | 0.25 | SERA | 0.13 |
| CRYM | 0.51 | MAPRE1 | 0.23 | ADD1 | 0.13 |
| CNP | 0.47 | LDHB | 0.21 | Atp6v0a1 | 0.13 |
| ALDOA | 0.43 | Atp6v1b2 | 0.20 | CPNE7 | 0.11 |
| GDA | 0.38 | PDXK | 0.20 | ARP3 | 0.11 |
| CSRP1 | 0.35 | PRRT2 | 0.19 | AGAP3 | 0.10 |
| RAB3A | 0.32 | Hsd17b4 | 0.18 | AMPH | 0.10 |
| CCT4 | 0.31 | RPH3A | 0.17 | SEC23A | 0.10 |

**Common partners with emPAI ratios WT/ LGI1 KO**

| | | | | | | | |
|---|---|---|---|---|---|---|---|
| MAP1B | 2.9 | AP3B2 | 1.57 | Phyhip | 0.54 | Csnk2a1 | 1.00 |
| MTAP2 | 2.7 | FAK1 | 1.53 | AP2A2 | 1.19 | SHPS1 | 0.96 |
| Rab11fip5 | 2.6 | ITIH2 | 1.50 | GIT1 | 1.11 | SYT2 | 0.95 |
| KCNA2 | 2.6 | Pip4k2b | 1.47 | AP2B1 | 1.10 | AP2A1 | 0.94 |
| Arhgap26 | 2.4 | Sh2d3c | 1.38 | Vtn | 1.10 | Lancl2 | 0.94 |
| VAPA | 2.3 | Epb41l3 (4.1B) | 1.38 | BCAS1 | 1.09 | SYT1 | 0.90 |
| MAP1A | 2.2 | Atp1a1 | 1.34 | CAMKV | 1.09 | BASP1 | 0.87 |
| Gramd1b | 2.1 | Atp1a3 | 1.32 | Cmas | 1.07 | Gap43 | 0.87 |
| JIP3 | 1.9 | KIF2A | 1.31 | MAP4 | 1.06 | Slc1a2 | 0.81 |
| Srcin1 | 1.8 | Camk2d | 1.28 | Ywhaq | 1.05 | DCX | 0.81 |
| PDCD6 | 1.8 | Atp6v1h | 1.27 | Iqsec1 | 1.04 | tcp1 | 0.78 |
| Camk2g | 1.7 | AP1B1 | 1.27 | KIF5C | 1.03 | CNTN1 | 0.76 |
| Atp2b1 | 1.7 | Map1lc3a | 1.26 | GSK3B | 1.00 | PRPS1 | 0.75 |
| Ppp3cb | 1.6 | cyfip2 | 1.25 | BCAR1 | 1.00 | AP2M1 | 0.73 |
| MAP6 | 1.6 | GRIA2 | 1.25 | SRC | 1.00 | GNAI1 | 0.50 |
| Thbs1 | 1.57 | CP | 1.22 | Pcmtd1 | 1.00 | Cct2 | 0.39 |
| EFS | 1.00 | Csnk2a2 | 0.70 | | | | |

Numbers represent emPAIs for specific partners and emPAIs ratios for common partners

### ESM_7
### Common partners of $pY_{458}Kv_{1.2}$ in WT or *Lgi1*$^{-/-}$

#### ESM_7: Common partners of $pY_{458}Kv_{1.2}$ in WT and Lgi1$^{-/-}$

| Common_IDs | emPAI_WT | emPAI_KO | emPAI_WT/emPAI_KO | log(emPAI_WT/emPAI_KO) | Description |
|---|---|---|---|---|---|
| **MAP1B** | 0.755 | 0.26 | **2.903846154** | 0.462973604 | Microtubule-associated protein 1B OS=Mus musculus OX=10090 GN=Map1b PE=1 SV=2 |
| **MTAP2** | 0.358333333 | 0.133333333 | **2.6875** | 0.429348473 | Microtubule-associated protein 2 OS=Mus musculus OX=10090 GN=Map2 PE=1 SV=2 |
| **RFIP5 (Rab11fip5)** | 0.26 | 0.1 | **2.6** | 0.414973348 | Rab11 family-interacting protein 5 OS=Mus musculus OX=10090 GN=Rab11fip5 PE=1 SV=2 |
| KCNA2 | 0.36 | 0.14 | **2.571428571** | 0.410174465 | **Potassium voltage-gated channel subfamily A member 2 OS=Mus musculus OX=10090 GN=Kcna2 PE=1 SV=1** |
| **RHG26 (Arhgap26)** | 0.263333333 | 0.11 | **2.393939394** | 0.379113151 | Rho GTPase-activating protein 26 OS=Mus musculus OX=10090 GN=Arhgap26 PE=1 SV=3 |
| **VAPA** | 0.57 | 0.25 | **2.28** | 0.357934847 | Vesicle-associated membrane protein-associated protein A OS=Mus musculus OX=10090 GN=Vapa PE=1 SV=2 |
| **MAP1A** | 0.133333333 | 0.06 | **2.222222222** | 0.346787486 | Microtubule-associated protein 1A OS=Mus musculus OX=10090 GN=Map1a PE=1 SV=2 |
| **ASTRB (Gramd1b)** | 0.316666667 | 0.15 | **2.111111111** | 0.324511092 | Protein Aster-B OS=Mus musculus OX=10090 GN=Gramd1b PE=1 SV=2 |
| **JIP3 (Mapk8ip3)** | 0.14 | 0.073333333 | **1.909090909** | 0.28082661 | C-Jun-amino-terminal kinase-interacting protein 3 OS=Mus musculus OX=10090 GN=Mapk8ip3 PE=1 SV=1 |
| **SRCN1 (Srcin1)** | 0.203333333 | 0.11 | **1.848484848** | 0.266815895 | SRC kinase signaling inhibitor 1 OS=Mus musculus OX=10090 GN=Srcin1 PE=1 SV=2 |
| **PDCD6** | 0.7 | 0.396666667 | **1.764705882** | 0.246672333 | Programmed cell death protein 6 OS=Mus musculus OX=10090 GN=Pdcd6 PE=1 SV=2 |
| **KCC2G (Camk2g)** | 0.725 | 0.43 | **1.686046512** | 0.226869551 | Calcium/calmodulin-dependent protein kinase type II subunit gamma OS=Mus musculus OX=10090 GN=Camk2g PE=1 SV=1 |
| **ATP2B1** | 0.19 | 0.113333333 | **1.676470588** | 0.224395939 | Plasma membrane calcium-transporting ATPase 1 OS=Mus musculus OX=10090 GN=Atp2b1 PE=1 SV=1 |
| **PP2BB (Ppp3cb)** | 0.41 | 0.255 | **1.607843137** | 0.206243676 | Serine/threonine-protein phosphatase 2B catalytic subunit beta isoform OS=Mus musculus OX=10090 GN=Ppp3cb PE=1 SV=2 |
| **MAP6** | 0.65 | 0.408333333 | **1.591836735** | 0.201898523 | Microtubule-associated protein 6 OS=Mus musculus OX=10090 GN=Map6 PE=1 SV=2 |
| **TSP1 (Thbs1)** | 0.22 | 0.14 | **1.571428571** | 0.196294645 | Thrombospondin-1 OS=Mus musculus OX=10090 GN=Thbs1 PE=1 SV=1 |
| **AP3B2** | 0.11 | 0.07 | **1.571428571** | 0.196294645 | AP-3 complex subunit beta-2 OS=Mus musculus OX=10090 GN=Ap3b2 PE=1 SV=2 |
| **FAK1 (Ptk2)** | 3.095 | 2.016666667 | **1.534710744** | 0.186026533 | Focal adhesion kinase 1 OS=Mus musculus OX=10090 GN=Ptk2 PE=1 SV=4 |
| **ITIH2** | 0.11 | 0.073333333 | **1.5** | 0.176091259 | Inter-alpha-trypsin inhibitor heavy chain H2 OS=Mus musculus OX=10090 GN=Itih2 PE=1 SV=1 |
| **PI42B (Pip4k2b)** | 0.193333333 | 0.131666667 | **1.46835443** | 0.166830898 | Phosphatidylinositol 5-phosphate 4-kinase type-2 beta OS=Mus musculus OX=10090 GN=Pip4k2b PE=1 SV=1 |
| **SH2D3 (Sh2d3c)** | 0.096666667 | 0.07 | **1.380952381** | 0.140178703 | SH2 domain-containing protein 3C OS=Mus musculus OX=10090 GN=Sh2d3c PE=1 SV=1 |
| **E41L3 (Epb41l3; 4.1B)** | 0.133333333 | 0.096666667 | **1.379310345** | 0.139661993 | Band 4.1-like protein 3 OS=Mus musculus OX=10090 GN=Epb41l3 PE=1 SV=1 |
| **ATP1A1** | 0.495 | 0.37 | **1.337837838** | 0.126403475 | Sodium/potassium-transporting ATPase subunit alpha-1 OS=Mus musculus OX=10090 GN=Atp1a1 PE=1 SV=1 |
| **ATP1A3** | 0.685 | 0.52 | **1.317307692** | 0.119687228 | Sodium/potassium-transporting ATPase subunit alpha-3 OS=Mus musculus OX=10090 GN=Atp1a3 PE=1 SV=1 |
| **KIF2A** | 0.213333333 | 0.163333333 | **1.306122449** | 0.115983894 | Kinesin-like protein KIF2A OS=Mus musculus OX=10090 GN=Kif2a PE=1 SV=2 |
| **KCC2D (Camk2d)** | 0.705 | 0.55 | **1.281818182** | 0.107826427 | Calcium/calmodulin-dependent protein kinase type II subunit delta OS=Mus musculus OX=10090 GN=Camk2d PE=1 SV=1 |
| **VATH (Atp6v1h)** | 0.165 | 0.13 | **1.269230769** | 0.103540592 | V-type proton ATPase subunit H OS=Mus musculus OX=10090 GN=Atp6v1h PE=1 SV=1 |
| **AP1B1** | 0.52 | 0.41 | **1.268292683** | 0.103219487 | AP-1 complex subunit beta-1 OS=Mus musculus OX=10090 GN=Ap1b1 PE=1 SV=2 |
| **MLP3A (Map1lc3a)** | 0.91 | 0.725 | **1.255172414** | 0.098703386 | Microtubule-associated proteins 1A/1B light chain 3A OS=Mus musculus OX=10090 GN=Map1lc3a PE=1 SV=1 |
| **CYFIP2** | 0.106666667 | 0.085 | **1.254901961** | 0.098609798 | Cytoplasmic FMR1-interacting protein 2 OS=Mus musculus OX=10090 GN=Cyfip2 PE=1 SV=2 |
| **GRIA2** | 0.106666667 | 0.085 | **1.254901961** | 0.098609798 | Glutamate receptor 2 OS=Mus musculus OX=10090 GN=Gria2 PE=1 SV=3 |
| **CERU (Cp)** | 0.11 | 0.09 | **1.222222222** | 0.087150176 | Ceruloplasmin OS=Mus musculus OX=10090 GN=Cp PE=1 SV=2 |
| **CLASP2** | 0.096666667 | 0.08 | **1.208333333** | 0.082186756 | CLIP-associating protein 2 OS=Mus musculus OX=10090 GN=Clasp2 PE=1 SV=1 |
| **AP2A2** | 0.746666667 | 0.63 | **1.185185185** | 0.073786214 | AP-2 complex subunit alpha-2 OS=Mus musculus OX=10090 GN=Ap2a2 PE=1 SV=2 |
| **GIT1** | 0.163333333 | 0.146666667 | **1.113636364** | 0.046743404 | ARF GTPase-activating protein GIT1 OS=Mus musculus OX=10090 GN=Git1 PE=1 SV=1 |
| **AP2B1** | 0.773333333 | 0.703333333 | **1.099526066** | 0.04120553 | AP-2 complex subunit beta OS=Mus musculus OX=10090 GN=Ap2b1 PE=1 SV=1 |
| **VTN** | 0.26 | 0.236666667 | **1.098591549** | 0.040836254 | Vitronectin OS=Mus musculus OX=10090 GN=Vtn PE=1 SV=2 |
| **BCAS1** | 0.145 | 0.133333333 | **1.0875** | 0.036429266 | Breast carcinoma-amplified sequence 1 homolog OS=Mus musculus OX=10090 GN=Bcas1 PE=1 SV=3 |
| **CAMKV** | 0.673333333 | 0.62 | **1.086021505** | 0.035838425 | CaM kinase-like vesicle-associated protein OS=Mus musculus OX=10090 GN=Camkv PE=1 SV=2 |
| **NEUA (Cmas)** | 0.193333333 | 0.18 | **1.074074074** | 0.031034234 | N-acylneuraminate cytidylyltransferase OS=Mus musculus OX=10090 GN=Cmas PE=1 SV=2 |
| **MAP4** | 0.206666667 | 0.195 | **1.05982906** | 0.025235823 | Microtubule-associated protein 4 OS=Mus musculus OX=10090 GN=Map4 PE=1 SV=3 |
| **1433T (Ywhaq)** | 1.45 | 1.375 | **1.054545455** | 0.023065304 | 14-3-3 protein theta OS=Mus musculus OX=10090 GN=Ywhaq PE=1 SV=1 |
| **NUCL** | 0.136666667 | 0.13 | **1.051282051** | 0.02171925 | Nucleolin OS=Mus musculus OX=10090 GN=Ncl PE=1 SV=2 |
| **KIF5C** | 0.113333333 | 0.11 | **1.03030303** | 0.012964977 | Kinesin heavy chain isoform 5C OS=Mus musculus OX=10090 GN=Kif5c PE=1 SV=3 |
| **GSK3B** | 0.195 | 0.195 | **1** | 9.64327E-17 | Glycogen synthase kinase-3 beta OS=Mus musculus OX=10090 GN=Gsk3b PE=1 SV=2 |
| **BCAR1** | 0.273333333 | 0.273333333 | **1** | 0 | Breast cancer anti-estrogen resistance protein 1 OS=Mus musculus OX=10090 GN=Bcar1 PE=1 SV=2 |
| **SRC** | 0.14 | 0.14 | **1** | 0 | Neuronal proto-oncogene tyrosine-protein kinase Src OS=Mus musculus OX=10090 GN=Src PE=1 SV=4 |
| **PCMTD1** | 0.13 | 0.13 | **1** | 0 | Protein-L-isoaspartate O-methyltransferase domain-containing protein 1 OS=Mus musculus OX=10090 GN=Pcmtd1 PE=1 SV=1 |
| **EFS** | 0.11 | 0.11 | **1** | 0 | Embryonal Fyn-associated substrate OS=Mus musculus OX=10090 GN=Efs PE=1 SV=2 |
| **DIP2B** | 0.04 | 0.04 | **1** | 0 | Disco-interacting protein 2 homolog B OS=Mus musculus OX=10090 GN=Dip2b PE=1 SV=1 |
| **CSK21 (Csnk2a1)** | 0.69 | 0.691666667 | **0.997590361** | -0.001047756 | Casein kinase II subunit alpha OS=Mus musculus OX=10090 GN=Csnk2a1 PE=1 SV=2 |
| **SHPS1 (Sirpa)** | 0.496666667 | 0.516666667 | **0.961290323** | -0.01714543 | Tyrosine-protein phosphatase non-receptor type substrate 1 OS=Mus musculus OX=10090 GN=Sirpa PE=1 SV=2 |
| **TNIK** | 0.06 | 0.063333333 | **0.947368421** | -0.023481096 | Traf2 and NCK-interacting protein kinase OS=Mus musculus OX=10090 GN=Tnik PE=1 SV=2 |
| **SYT2** | 0.753333333 | 0.796666667 | **0.945606695** | -0.024289462 | Synaptotagmin-2 OS=Mus musculus OX=10090 GN=Syt2 PE=1 SV=1 |
| **AP2A1** | 0.493333333 | 0.523333333 | **0.942675159** | -0.025637937 | AP-2 complex subunit alpha-1 OS=Mus musculus OX=10090 GN=Ap2a1 PE=1 SV=1 |
| **LANC2 (Lancl2)** | 0.493333333 | 0.523333333 | **0.942675159** | -0.025637937 | LanC-like protein 2 OS=Mus musculus OX=10090 GN=Lancl2 PE=1 SV=1 |

| | | | | | | |
|---|---|---|---|---|---|---|
| **SYT1** | 2.716666667 | 3.008333333 | **0.903047091** | | -0.044289602 | Synaptotagmin-1 OS=Mus musculus OX=10090 GN=Syt1 PE=1 SV=1 |
| **BASP1** | 0.69 | 0.79 | **0.873417722** | | -0.058778001 | Brain acid soluble protein 1 OS=Mus musculus OX=10090 GN=Basp1 PE=1 SV=3 |
| **NEUM (Gap43)** | 1.87 | 2.16 | **0.865740741** | | -0.062612145 | Neuromodulin OS=Mus musculus OX=10090 GN=Gap43 PE=1 SV=1 |
| **RTN4** | 0.065 | 0.08 | **0.8125** | | -0.09017663 | Reticulon-4 OS=Mus musculus OX=10090 GN=Rtn4 PE=1 SV=2 |
| **EAA2 (Slc1a2)** | 0.17 | 0.21 | **0.80952381** | | -0.091770373 | Excitatory amino acid transporter 2 OS=Mus musculus OX=10090 GN=Slc1a2 PE=1 SV=1 |
| **DCX** | 0.23 | 0.285 | **0.807017544** | | -0.093117024 | Neuronal migration protein doublecortin OS=Mus musculus OX=10090 GN=Dcx PE=1 SV=1 |
| **TCP1** | 0.14 | 0.18 | **0.777777778** | | -0.109144469 | T-complex protein 1 subunit alpha OS=Mus musculus OX=10090 GN=Tcp1 PE=1 SV=3 |
| **CAPR1** | 0.07 | 0.09 | **0.777777778** | | -0.109144469 | Caprin-1 OS=Mus musculus OX=10090 GN=Caprin1 PE=1 SV=2 |
| **CNTN1** | 0.118333333 | 0.155 | **0.76344086** | | -0.1172246 | Contactin-1 OS=Mus musculus OX=10090 GN=Cntn1 PE=1 SV=1 |
| **PRPS1** | 0.15 | 0.2 | **0.75** | | -0.124938737 | Ribose-phosphate pyrophosphokinase 1 OS=Mus musculus OX=10090 GN=Prps1 PE=1 SV=4 |
| **AP2M1** | 0.44 | 0.606666667 | **0.725274725** | | -0.139497457 | AP-2 complex subunit mu OS=Mus musculus OX=10090 GN=Ap2m1 PE=1 SV=1 |
| **FAS** | 0.031666667 | 0.045 | **0.703703704** | | -0.152610163 | Fatty acid synthase OS=Mus musculus OX=10090 GN=Fasn PE=1 SV=2 |
| **CSK22 (Csnk2a2)** | 0.363333333 | 0.516666667 | **0.703225806** | | -0.1529052 | Casein kinase II subunit alpha' OS=Mus musculus OX=10090 GN=Csnk2a2 PE=1 SV=1 |
| **PHYHIP** | 0.208333333 | 0.385 | **0.541125541** | | -0.266701967 | Phytanoyl-CoA hydroxylase-interacting protein OS=Mus musculus OX=10090 GN=Phyhip PE=1 SV=1 |
| **GNAI1** | 0.17 | 0.336666667 | **0.504950495** | | -0.296751198 | Guanine nucleotide-binding protein G(i) subunit alpha-1 OS=Mus musculus OX=10090 GN=Gnai1 PE=1 SV=1 |
| **TCPB (Cct2)** | 0.11 | 0.285 | **0.385964912** | | -0.413452175 | T-complex protein 1 subunit beta OS=Mus musculus OX=10090 GN=Cct2 PE=1 SV=4 |

### ESM_8
### Exclusive partners of $pY_{458}Kv_{1.2}$ in WT

#### ESM_8: Exclusive partners of $pY_{458}Kv_{1.2}$ in WT

| WT_exc_List | emPAI filtered | emPAI StdDev filtered | N times | StdError emPAI filtered | Description |
|---|---|---|---|---|---|
| PIN1_MOUSE | 1.23 | 0 | 2 | 0 | Peptidyl-prolyl cis-trans isomerase NIMA-interacting 1 OS=Mus musculus OX=10090 GN=Pin1 PE=1 SV=1 |
| NEUG_MOUSE | 1.18 | 0 | 2 | 0 | Neurogranin OS=Mus musculus OX=10090 GN=Nrgn PE=1 SV=1 |
| CYCS_MOUSE | 1.04 | 0 | 2 | 0 | Cytochrome c, somatic OS=Mus musculus OX=10090 GN=Cycs PE=1 SV=2 |
| GNAO_MOUSE | 0.903333333 | 0.300924501 | 3 | 0.173738842 | Guanine nucleotide-binding protein G(o) subunit alpha OS=Mus musculus OX=10090 GN=Gnao1 PE=1 SV=3 |
| 1433E_MOUSE | 0.78 | 0.11 | 2 | 0.077781746 | 14-3-3 protein epsilon OS=Mus musculus OX=10090 GN=Ywhae PE=1 SV=1 |
| TAU_MOUSE | 0.625 | 0.075 | 2 | 0.053033009 | Microtubule-associated protein tau OS=Mus musculus OX=10090 GN=Mapt PE=1 SV=3 |
| DYN1_MOUSE | 0.6 | 0.03 | 2 | 0.021213203 | Dynamin-1 OS=Mus musculus OX=10090 GN=Dnm1 PE=1 SV=2 |
| DIRA2_MOUSE | 0.505 | 0 | 2 | 0 | GTP-binding protein Di-Ras2 OS=Mus musculus OX=10090 GN=Diras2 PE=1 SV=1 |
| DCLK1_MOUSE | 0.455 | 0.15 | 2 | 0.106066017 | Serine/threonine-protein kinase DCLK1 OS=Mus musculus OX=10090 GN=Dclk1 PE=1 SV=1 |
| OTUB1_MOUSE | 0.425 | 0.075 | 2 | 0.053033009 | Ubiquitin thioesterase OTUB1 OS=Mus musculus OX=10090 GN=Otub1 PE=1 SV=2 |
| PP2BA_MOUSE | 0.41 | 0.04 | 2 | 0.028284271 | Serine/threonine-protein phosphatase 2B catalytic subunit alpha isoform OS=Mus musculus OX=10090 GN=Ppp3ca PE=1 SV=1 |
| GBB2_MOUSE | 0.39 | 0 | 2 | 0 | Guanine nucleotide-binding protein G(I)/G(S)/G(T) subunit beta-2 OS=Mus musculus OX=10090 GN=Gnb2 PE=1 SV=3 |
| BAIP2_MOUSE | 0.375 | 0.08 | 2 | 0.056568542 | Brain-specific angiogenesis inhibitor 1-associated protein 2 OS=Mus musculus OX=10090 GN=Baiap2 PE=1 SV=2 |
| CSK2B_MOUSE | 0.346666667 | 0.080138769 | 3 | 0.04626814 | Casein kinase II subunit beta OS=Mus musculus OX=10090 GN=Csnk2b PE=1 SV=1 |
| SPTN1_MOUSE (SPTAN1) | 0.315 | 0.055 | 2 | 0.038890873 | Spectrin alpha chain, non-erythrocytic 1 OS=Mus musculus OX=10090 GN=Sptan1 PE=1 SV=4 |
| COF1_MOUSE | 0.31 | 0 | 2 | 0 | Cofilin-1 OS=Mus musculus OX=10090 GN=Cfl1 PE=1 SV=3 |
| MK03_MOUSE | 0.295 | 0.045 | 2 | 0.031819805 | Mitogen-activated protein kinase 3 OS=Mus musculus OX=10090 GN=Mapk3 PE=1 SV=5 |
| CDK5_MOUSE | 0.29 | 0.065 | 2 | 0.045961941 | Cyclin-dependent-like kinase 5 OS=Mus musculus OX=10090 GN=Cdk5 PE=1 SV=1 |
| SC22B_MOUSE | 0.29 | 0 | 2 | 0 | Vesicle-trafficking protein SEC22b OS=Mus musculus OX=10090 GN=Sec22b PE=1 SV=3 |
| PDIA3_MOUSE | 0.28 | 0 | 2 | 0 | Protein disulfide-isomerase A3 OS=Mus musculus OX=10090 GN=Pdia3 PE=1 SV=2 |
| MRP_MOUSE | 0.275 | 0 | 2 | 0 | MARCKS-related protein OS=Mus musculus OX=10090 GN=Marcksl1 PE=1 SV=2 |
| PFKAM_MOUSE | 0.27 | 0.121928941 | 3 | 0.070395707 | ATP-dependent 6-phosphofructokinase, muscle type OS=Mus musculus OX=10090 GN=Pfkm PE=1 SV=3 |
| KCNA1_MOUSE | 0.265 | 0.145 | 2 | 0.102530483 | Potassium voltage-gated channel subfamily A member 1 OS=Mus musculus OX=10090 GN=Kcna1 PE=1 SV=1 |
| SNP25_MOUSE | 0.24 | 0 | 2 | 0 | Synaptosomal-associated protein 25 OS=Mus musculus OX=10090 GN=Snap25 PE=1 SV=1 |
| RACK1_MOUSE | 0.22 | 0 | 2 | 0 | Receptor of activated protein C kinase 1 OS=Mus musculus OX=10090 GN=Rack1 PE=1 SV=3 |
| BACH_MOUSE | 0.213333333 | 0.077602978 | 3 | 0.0448041 | Cytosolic acyl coenzyme A thioester hydrolase OS=Mus musculus OX=10090 GN=Acot7 PE=1 SV=2 |
| LDHA_MOUSE | 0.21 | 0 | 2 | 0 | L-lactate dehydrogenase A chain OS=Mus musculus OX=10090 GN=Ldha PE=1 SV=3 |
| SIRT2_MOUSE | 0.19 | 0.042426407 | 3 | 0.024494897 | NAD-dependent protein deacetylase sirtuin-2 OS=Mus musculus OX=10090 GN=Sirt2 PE=1 SV=2 |
| KCD12_MOUSE | 0.19 | 0 | 2 | 0 | BTB/POZ domain-containing protein KCTD12 OS=Mus musculus OX=10090 GN=Kctd12 PE=1 SV=1 |
| NONO_MOUSE | 0.19 | 0 | 2 | 0 | Non-POU domain-containing octamer-binding protein OS=Mus musculus OX=10090 GN=Nono PE=1 SV=3 |
| VPS35_MOUSE | 0.19 | 0.04 | 2 | 0.028284271 | Vacuolar protein sorting-associated protein 35 OS=Mus musculus OX=10090 GN=Vps35 PE=1 SV=1 |
| PON1_MOUSE | 0.17 | 0 | 3 | 0 | Serum paraoxonase/arylesterase 1 OS=Mus musculus OX=10090 GN=Pon1 PE=1 SV=2 |
| 2AAA_MOUSE | 0.17 | 0 | 2 | 0 | Serine/threonine-protein phosphatase 2A 65 kDa regulatory subunit A alpha isoform OS=Mus musculus OX=10090 GN=Ppp2r1a PE=1 S |
| GNB5_MOUSE | 0.16 | 0 | 2 | 0 | Guanine nucleotide-binding protein subunit beta-5 OS=Mus musculus OX=10090 GN=Gnb5 PE=1 SV=1 |
| KCAB2_MOUSE | 0.16 | 0.042426407 | 3 | 0.024494897 | Voltage-gated potassium channel subunit beta-2 OS=Mus musculus OX=10090 GN=Kcnab2 PE=1 SV=1 |
| SEPT5_MOUSE | 0.16 | 0 | 2 | 0 | Septin-5 OS=Mus musculus OX=10090 GN=Sept5 PE=1 SV=2 |
| BTBDH_MOUSE | 0.153333333 | 0.032998316 | 3 | 0.019051587 | BTB/POZ domain-containing protein 17 OS=Mus musculus OX=10090 GN=Btbd17 PE=1 SV=1 |
| RFIP2_MOUSE | 0.15 | 0.03 | 2 | 0.021213203 | Rab11 family-interacting protein 2 OS=Mus musculus OX=10090 GN=Rab11fip2 PE=1 SV=1 |
| PGK1_MOUSE | 0.145 | 0.042426407 | 3 | 0.024494897 | Phosphoglycerate kinase 1 OS=Mus musculus OX=10090 GN=Pgk1 PE=1 SV=4 |
| ARHG2_MOUSE | 0.143333333 | 0.065489609 | 3 | 0.037810443 | Rho guanine nucleotide exchange factor 2 OS=Mus musculus OX=10090 GN=Arhgef2 PE=1 SV=4 |
| SEP11_MOUSE | 0.14 | 0 | 3 | 0 | Septin-11 OS=Mus musculus OX=10090 GN=Sept11 PE=1 SV=4 |
| SEPT8_MOUSE | 0.14 | 0 | 2 | 0 | Septin-8 OS=Mus musculus OX=10090 GN=Sept8 PE=1 SV=4 |
| WASF1_MOUSE | 0.14 | 0.03 | 2 | 0.021213203 | Wiskott-Aldrich syndrome protein family member 1 OS=Mus musculus OX=10090 GN=Wasf1 PE=1 SV=2 |
| ABI1_MOUSE | 0.135 | 0.035 | 2 | 0.024748737 | Abl interactor 1 OS=Mus musculus OX=10090 GN=Abi1 PE=1 SV=3 |
| COR1C_MOUSE | 0.13 | 0 | 2 | 0 | Coronin-1C OS=Mus musculus OX=10090 GN=Coro1c PE=1 SV=2 |
| SPTB2_MOUSE(SPTBN1) | 0.123333333 | 0.012472191 | 3 | 0.007200823 | Spectrin beta chain, non-erythrocytic 1 OS=Mus musculus OX=10090 GN=Sptbn1 PE=1 SV=2 |
| ASAP1_MOUSE | 0.11 | 0.024494897 | 3 | 0.014142136 | Arf-GAP with SH3 domain, ANK repeat and PH domain-containing protein 1 OS=Mus musculus OX=10090 GN=Asap1 PE=1 SV=2 |
| EAA1_MOUSE (Slc1a3) | 0.11 | 0 | 2 | 0 | Excitatory amino acid transporter 1 OS=Mus musculus OX=10090 GN=Slc1a3 PE=1 SV=2 |
| G3BP2_MOUSE | 0.1 | 0 | 2 | 0 | Ras GTPase-activating protein-binding protein 2 OS=Mus musculus OX=10090 GN=G3bp2 PE=1 SV=2 |
| PFKAL_MOUSE | 0.1 | 0.02 | 2 | 0.014142136 | ATP-dependent 6-phosphofructokinase, liver type OS=Mus musculus OX=10090 GN=Pfkl PE=1 SV=4 |

### ESM_9
### Exclusive partners of $pY_{458}Kv_{1.2}$ in *Lgi1*$^{-/-}$

| ESM_9: Exclusive partners of $pY_{458}Kv_{1.2}$ in Lgi1$^{-/-}$ | | | | | |
|---|---|---|---|---|---|
| KO_exc_list | emPAI filtered | emPAI StdDev filtered | N times | StdError emPAI filtered | Description |
| G3P_MOUSE | 1.525 | 0 | 2 | 0 | Glyceraldehyde-3-phosphate dehydrogenase OS=Mus musculus OX=19 GN=Gapdh PE=1 SV=2 |
| MT3_MOUSE | 1.025 | 0 | 3 | 0 | Metallothionein-3 OS=Mus musculus OX=19 GN=Mt3 PE=1 SV=1 |
| RAC1_MOUSE | 0.93 | 0.14 | 2 | 0.098994949 | Ras-related C3 botulinum toxin substrate 1 OS=Mus musculus OX=19 GN=Rac1 PE=1 SV=1 |
| ATP1B1_MOUSE | 0.9 | 0.095 | 2 | 0.067175144 | Sodium/potassium-transporting ATPase subunit beta-1 OS=Mus musculus OX=19 GN=Atp1b1 PE=1 SV=1 |
| RAC3_MOUSE | 0.815 | 0.265 | 2 | 0.187383297 | Ras-related C3 botulinum toxin substrate 3 OS=Mus musculus OX=19 GN=Rac3 PE=1 SV=1 |
| MARCS_MOUSE | 0.723333333 | 0.103708995 | 3 | 0.059876416 | Myristoylated alanine-rich C-kinase substrate OS=Mus musculus OX=19 GN=Marcks PE=1 SV=2 |
| THIO_MOUSE (TXN) | 0.68 | 0 | 2 | 0 | Thioredoxin OS=Mus musculus OX=19 GN=Txn PE=1 SV=3 |
| CRYM_MOUSE | 0.511666667 | 0.202703944 | 3 | 0.117031177 | Ketimine reductase mu-crystallin OS=Mus musculus OX=19 GN=Crym PE=1 SV=1 |
| CN37_MOUSE | 0.465 | 0.133666251 | 3 | 0.077172246 | 2',3'-cyclic-nucleotide 3'-phosphodiesterase OS=Mus musculus OX=19 GN=Cnp PE=1 SV=3 |
| ALDOA_MOUSE | 0.43 | 0.06 | 2 | 0.042426407 | Fructose-bisphosphate aldolase A OS=Mus musculus OX=19 GN=Aldoa PE=1 SV=2 |
| GUAD_MOUSE | 0.375 | 0.045 | 2 | 0.031819805 | Guanine deaminase OS=Mus musculus OX=19 GN=Gda PE=1 SV=1 |
| RAB3A_MOUSE | 0.315 | 0 | 3 | 0 | Ras-related protein Rab-3A OS=Mus musculus OX=19 GN=Rab3a PE=1 SV=1 |
| TCPD_MOUSE (Cct4) | 0.31 | 0.04 | 2 | 0.028284271 | T-complex protein 1 subunit delta OS=Mus musculus OX=19 GN=Cct4 PE=1 SV=3 |
| FHL1_MOUSE | 0.285 | 0.065 | 2 | 0.045961941 | Four and a half LIM domains protein 1 OS=Mus musculus OX=19 GN=Fhl1 PE=1 SV=3 |
| STX1B_MOUSE | 0.27 | 0.06 | 2 | 0.042426407 | Syntaxin-1B OS=Mus musculus OX=19 GN=Stx1b PE=1 SV=1 |
| SYPH_MOUSE | 0.27 | 0 | 2 | 0 | Synaptophysin OS=Mus musculus OX=19 GN=Syp PE=1 SV=2 |
| GAS7_MOUSE | 0.25 | 0.104243305 | 3 | 0.0601849 | Growth arrest-specific protein 7 OS=Mus musculus OX=19 GN=Gas7 PE=1 SV=1 |
| KPCG_MOUSE | 0.246666667 | 0.089566859 | 3 | 0.05171145 | Protein kinase C gamma type OS=Mus musculus OX=19 GN=Prkcg PE=1 SV=1 |
| FA49B_MOUSE (CYRIB) | 0.245 | 0.055 | 2 | 0.038890873 | Protein FAM49B OS=Mus musculus OX=19 GN=Fam49b PE=1 SV=1 |
| MAPRE1_MOUSE | 0.23 | 0 | 3 | 0 | Microtubule-associated protein RP/EB family member 1 OS=Mus musculus OX=19 GN=Mapre1 PE=1 SV=3 |
| LDHB_MOUSE | 0.21 | 0 | 2 | 0 | L-lactate dehydrogenase B chain OS=Mus musculus OX=19 GN=Ldhb PE=1 SV=2 |
| VATB2_MOUSE (Atp6v1b2) | 0.196666667 | 0.057348835 | 3 | 0.033110365 | V-type proton ATPase subunit B, brain isoform OS=Mus musculus OX=19 GN=Atp6v1b2 PE=1 SV=1 |
| PDXK_MOUSE | 0.191666667 | 0.051854497 | 3 | 0.029938208 | Pyridoxal kinase OS=Mus musculus OX=19 GN=Pdxk PE=1 SV=1 |
| PRRT2_MOUSE | 0.19 | 0 | 2 | 0 | Proline-rich transmembrane protein 2 OS=Mus musculus OX=19 GN=Prrt2 PE=1 SV=1 |
| DHB4_MOUSE (Hsd17b4 ) | 0.18 | 0.07 | 2 | 0.049497475 | Peroxisomal multifunctional enzyme type 2 OS=Mus musculus OX=19 GN=Hsd17b4 PE=1 SV=3 |
| RPH3A_MOUSE | 0.166666667 | 0.047140452 | 3 | 0.027216553 | Rabphilin-3A OS=Mus musculus OX=19 GN=Rph3a PE=1 SV=2 |
| ACTZ_MOUSE (Actr1a ) | 0.16 | 0 | 2 | 0 | Alpha-centractin OS=Mus musculus OX=19 GN=Actr1a PE=1 SV=1 |
| KAP3_MOUSE | 0.16 | 0 | 2 | 0 | cAMP-dependent protein kinase type II-beta regulatory subunit OS=Mus musculus OX=19 GN=Prkar2b PE=1 SV=3 |
| PFKP_MOUSE | 0.156666667 | 0.023570226 | 3 | 0.013608276 | ATP-dependent 6-phosphofructokinase, platelet type OS=Mus musculus OX=19 GN=Pfkp PE=1 SV=1 |
| UBA1_MOUSE | 0.15 | 0.03 | 2 | 0.021213203 | Ubiquitin-like modifier-activating enzyme 1 OS=Mus musculus OX=19 GN=Uba1 PE=1 SV=1 |
| TERA_MOUSE (VCP) | 0.136666667 | 0.03681787 | 3 | 0.021256807 | Transitional endoplasmic reticulum ATPase OS=Mus musculus OX=19 GN=Vcp PE=1 SV=4 |
| SEPTIN3_MOUSE | 0.13 | 0 | 2 | 0 | Neuronal-specific septin-3 OS=Mus musculus OX=19 GN=Sept3 PE=1 SV=2 |
| SERA_MOUSE | 0.13 | 0 | 2 | 0 | D-3-phosphoglycerate dehydrogenase OS=Mus musculus OX=19 GN=Phgdh PE=1 SV=3 |
| ADD1_MOUSE | 0.125 | 0.045 | 2 | 0.031819805 | Alpha-adducin OS=Mus musculus OX=19 GN=Add1 PE=1 SV=2 |
| VPP1_MOUSE (Atp6v0a1) | 0.125 | 0 | 2 | 0 | V-type proton ATPase 116 kDa subunit a isoform 1 OS=Mus musculus OX=19 GN=Atp6va1 PE=1 SV=3 |
| CPNE7_MOUSE | 0.11 | 0 | 2 | 0 | Copine-7 OS=Mus musculus OX=19 GN=Cpne7 PE=1 SV=1 |
| ARP3_MOUSE | 0.105 | 0 | 2 | 0 | Actin-related protein 3 OS=Mus musculus OX=19 GN=Actr3 PE=1 SV=3 |
| AGAP3_MOUSE | 0.1 | 0 | 2 | 0 | Arf-GAP with GTPase, ANK repeat and PH domain-containing protein 3 OS=Mus musculus OX=19 GN=Agap3 PE=1 SV=1 |
| AMPH_MOUSE | 0.1 | 0 | 2 | 0 | Amphiphysin OS=Mus musculus OX=19 GN=Amph PE=1 SV=1 |
| SEC23A_MOUSE | 0.1 | 0.04 | 2 | 0.028284271 | Protein transport protein Sec23A OS=Mus musculus OX=19 GN=Sec23a PE=1 SV=2 |

### ESM_S10

#### ESM_10
#### GO terms of $pY_{458}Kv_{1.2}$ partners in WT and *Lgi1*$^{-/-}$

##### GO terms subcellular localisation for $pY_{458}Kv_{1.2}$ partners in WT & $pY_{458}Kv_{1.2}$ partners in KO

###### Lost GO for $pY_{458}Kv_{1.2}$ in *Lgi1*$^{-/-}$

| GO term | Description | Partner NB |
|---|---|---|
| GOCC:0005622 | Intracellular | 45 |
| GOCC:0005737 | Cytoplasm | 42 |
| GOCC:0032991 | Protein-containing complex | 34 |
| GOCC:0030054 | Cell junction | 29 |
| GOCC:0120025 | Plasma membrane bounded cell projection | 28 |
| GOCC:0045202 | Synapse | 26 |
| GOCC:0036477 | Somatodendritic compartment | 15 |
| GOCC:0098796 | Membrane protein complex | 14 |
| GOCC:0044297 | Cell body | 11 |
| GOCC:0098984 | Neuron to neuron synapse | 11 |
| GOCC:0099572 | Postsynaptic specialization | 11 |
| GOCC:0005739 | Mitochondrion | 10 |
| GOCC:0014069 | Postsynaptic density | 10 |
| GOCC:0031252 | Cell leading edge | 10 |
| GOCC:0043025 | Neuronal cell body | 10 |
| GOCC:0098797 | Plasma membrane protein complex | 8 |
| GOCC:0030027 | Lamellipodium | 7 |
| GOCC:0044304 | Main axon | 7 |
| GOCC:0005938 | Cell cortex | 6 |
| GOCC:0030426 | Growth cone | 6 |
| GOCC:0098978 | Glutamatergic synapse | 6 |
| GOCC:0030175 | Filopodium | 5 |
| GOCC:0031234 | Extrinsic component of cytoplasmic side of plasma membrane | 5 |
| GOCC:0045121 | Membrane raft | 5 |
| GOCC:0044306 | Neuron projection terminus | 4 |
| GOCC:0005834 | Heterotrimeric G-protein complex | 3 |
| GOCC:0005940 | Septin ring | 3 |
| GOCC:0008076 | Voltage-gated potassium channel complex | 3 |
| GOCC:0016328 | Lateral plasma membrane | 3 |
| GOCC:0030864 | Cortical actin cytoskeleton | 3 |
| GOCC:0031105 | Septin complex | 3 |
| GOCC:0032432 | Actin filament bundle | 3 |
| GOCC:0033270 | Paranode region of axon | 3 |
| GOCC:0044224 | Juxtaparanode region of axon | 3 |
| GOCC:0098685 | Schaffer collateral - CA1 synapse | 3 |
| GOCC:0099522 | Region of cytosol | 3 |
| GOCC:0001891 | Phagocytic cup | 2 |
| GOCC:0005945 | 6-phosphofructokinase complex | 2 |
| GOCC:0005963 | Magnesium-dependent protein serine/threonine phosphatase complex | 2 |
| GOCC:0008091 | Spectrin | 2 |
| GOCC:0030673 | Axolemma | 2 |
| GOCC:0031209 | SCAR complex | 2 |
| GOCC:0032437 | Cuticular plate | 2 |
| GOCC:0033010 | Paranodal junction | 2 |
| GOCC:0043194 | Axon initial segment | 2 |
| GOCC:0044305 | Calyx of Held | 2 |
| GOCC:0097470 | Ribbon synapse | 2 |
| GOCC:0099026 | Anchored component of presynaptic membrane | 2 |
| GOCC:1990350 | Glucose transporter complex | 2 |
| GOCC:0099524 | Postsynaptic cytosol | 2 |

###### Gained GO for $pY_{458}Kv_{1.2}$ in *Lgi1*$^{-/-}$

| GO term | Description | Partner NB |
|---|---|---|
| GOCC:0012505 | Endomembrane system | 15 |
| GOCC:0031982 | Vesicle | 14 |
| GOCC:0031090 | Organelle membrane | 13 |
| GOCC:0031410 | Cytoplasmic vesicle | 13 |
| GOCC:0098588 | Bounding membrane of organelle | 10 |
| GOCC:0099513 | Polymeric cytoskeletal fiber | 10 |
| GOCC:0030659 | Cytoplasmic vesicle membrane | 8 |
| GOCC:0099503 | Secretory vesicle | 8 |
| GOCC:0005929 | Cilium | 6 |
| GOCC:0030658 | Transport vesicle membrane | 6 |
| GOCC:0043679 | Axon terminus | 6 |
| GOCC:0044309 | Neuron spine | 6 |
| GOCC:0030672 | Synaptic vesicle membrane | 5 |
| GOCC:0030135 | Coated vesicle | 4 |
| GOCC:0048471 | Perinuclear region of cytoplasm | 4 |
| GOCC:0005884 | Actin filament | 3 |
| GOCC:0030662 | Coated vesicle membrane | 3 |
| GOCC:0031594 | Neuromuscular junction | 3 |
| GOCC:0048786 | Presynaptic active zone | 3 |
| GOCC:0098563 | Intrinsic component of synaptic vesicle membrane | 3 |
| GOCC:0016471 | Vacuolar proton-transporting V-type ATPase complex | 2 |
| GOCC:0060203 | Clathrin-sculpted glutamate transport vesicle membrane | 2 |
| GOCC:0098850 | Extrinsic component of synaptic vesicle membrane | 2 |

###### Common GO for $pY_{458}Kv_{1.2}$ in WT & *Lgi1*$^{-/-}$

| GO term | Description | Partner NB |
|---|---|---|
| GOCC:0110165 | Cellular anatomical entity | 47 |
| GOCC:0110165 | Cellular anatomical entity | 41 |
| GOCC:0043226 | Organelle | 41 |
| GOCC:0043226 | Organelle | 35 |
| GOCC:0043229 | Intracellular organelle | 38 |
| GOCC:0043229 | Intracellular organelle | 34 |
| GOCC:0016020 | Membrane | 31 |
| GOCC:0016020 | Membrane | 24 |
| GOCC:0043227 | Membrane-bounded organelle | 31 |
| GOCC:0043227 | Membrane-bounded organelle | 31 |
| GOCC:0071944 | Cell periphery | 27 |
| GOCC:0071944 | Cell periphery | 19 |
| GOCC:0043231 | Intracellular membrane-bounded organelle | 28 |
| GOCC:0043231 | Intracellular membrane-bounded organelle | 28 |
| GOCC:0005829 | Cytosol | 26 |
| GOCC:0005829 | Cytosol | 18 |
| GOCC:0005886 | Plasma membrane | 25 |
| GOCC:0005886 | Plasma membrane | 17 |
| GOCC:0043232 | Intracellular non-membrane-bounded organelle | 22 |
| GOCC:0043232 | Intracellular non-membrane-bounded organelle | 17 |
| GOCC:0043005 | Neuron projection | 22 |
| GOCC:0043005 | Neuron projection | 13 |
| GOCC:0098794 | Postsynapse | 20 |
| GOCC:0098794 | Postsynapse | 9 |
| GOCC:0005856 | Cytoskeleton | 18 |
| GOCC:0005856 | Cytoskeleton | 15 |
| GOCC:0098590 | Plasma membrane region | 15 |
| GOCC:0098590 | Plasma membrane region | 12 |
| GOCC:0098793 | Presynapse | 12 |
| GOCC:0098793 | Presynapse | 11 |
| GOCC:0099080 | Supramolecular complex | 12 |
| GOCC:0099080 | Supramolecular complex | 10 |
| GOCC:0099512 | Supramolecular fiber | 11 |
| GOCC:0099512 | Supramolecular fiber | 9 |
| GOCC:0030424 | Axon | 13 |
| GOCC:0030424 | Axon | 8 |
| GOCC:0030425 | Dendrite | 12 |
| GOCC:0030425 | Dendrite | 6 |
| GOCC:0015629 | Actin cytoskeleton | 10 |
| GOCC:0015629 | Actin cytoskeleton | 7 |
| GOCC:0015630 | Microtubule cytoskeleton | 9 |
| GOCC:0015630 | Microtubule cytoskeleton | 9 |
| GOCC:0150034 | Distal axon | 9 |
| GOCC:0150034 | Distal axon | 7 |
| GOCC:0030133 | Transport vesicle | 8 |
| GOCC:0030133 | Transport vesicle | 5 |
| GOCC:0097060 | Synaptic membrane | 8 |
| GOCC:0097060 | Synaptic membrane | 7 |
| GOCC:0008021 | Synaptic vesicle | 7 |
| GOCC:0008021 | Synaptic vesicle | 4 |
| GOCC:0043197 | Dendritic spine | 6 |
| GOCC:0043197 | Dendritic spine | 5 |
| GOCC:0005874 | Microtubule | 6 |
| GOCC:0005874 | Microtubule | 6 |
| GOCC:0043209 | Myelin sheath | 6 |
| GOCC:0043209 | Myelin sheath | 4 |
| GOCC:0042734 | Presynaptic membrane | 5 |
| GOCC:0042734 | Presynaptic membrane | 5 |
| GOCC:0099523 | Presynaptic cytosol | 2 |
| GOCC:0099523 | Presynaptic cytosol | 2 |
